# Ca^2+^ channel and active zone protein abundance intersects with input-specific synapse organization to shape functional synaptic diversity

**DOI:** 10.1101/2023.04.02.535290

**Authors:** A. T. Medeiros, S.J. Gratz, A. Delgado, J.T. Ritt, Kate M. O’Connor-Giles

**Author notes:** For correspondence: Scott J. Gratz 185 Meeting St., Providence, RI 02912.; Kate M. O’Connor-Giles, 185 Meeting St., Providence, RI 02912. The authors declare no competing interests.

## Abstract

Synaptic heterogeneity is a hallmark of nervous systems that enables complex and adaptable communication in neural circuits. To understand circuit function, it is thus critical to determine the factors that contribute to the functional diversity of synapses. We investigated the contributions of voltage-gated calcium channel (VGCC) abundance, spatial organization, and subunit composition to synapse diversity among and between synapses formed by two closely related *Drosophila* glutamatergic motor neurons with distinct neurotransmitter release probabilities (P_r_). Surprisingly, VGCC levels are highly predictive of heterogeneous P_r_ among individual synapses of either low- or high-P_r_ inputs, but not between inputs. We find that the same number of VGCCs are more densely organized at high-P_r_ synapses, consistent with tighter VGCC-synaptic vesicle coupling. We generated endogenously tagged lines to investigate VGCC subunits *in vivo* and found that the α2δ-3 subunit Straightjacket along with the CAST/ELKS active zone (AZ) protein Bruchpilot, both key regulators of VGCCs, are less abundant at high-P_r_ inputs, yet positively correlate with P_r_ among synapses formed by either input. Consistently, both Straightjacket and Bruchpilot levels are dynamically increased across AZs of both inputs when neurotransmitter release is potentiated to maintain stable communication following glutamate receptor inhibition. Together, these findings suggest a model in which VGCC and AZ protein abundance intersects with input-specific spatial and molecular organization to shape the functional diversity of synapses.

## INTRODUCTION

The broad and complex functions of neural circuits depend on diverse neuronal subtypes communicating through synapses with distinct properties. Thus, understanding how synaptic diversity is established is critical for understanding neural circuit function. Neurotransmission occurs at specialized membranes called active zones (AZs) where action potentials drive the opening of voltage-gated Ca^2+^ channels (VGCCs) to trigger Ca^2+^-dependent synaptic vesicle (SV) fusion and neurotransmitter release. Neurotransmitter release properties are determined locally at individual synapses and vary considerably between neuronal subtypes and within homogeneous populations of neurons (Ariel, Hoppa, and Ryan 2012; Atwood and Karunanithi 2002; Branco and Staras 2009; Hatt and Smith 1976). In fact, functional imaging studies in *Drosophila* demonstrate that even single neurons forming synapses with the same postsynaptic partner display heterogeneous synaptic strength among individual AZs (Guerrero et al. 2005; Melom et al. 2013; Peled and Isacoff 2011).

Presynaptic strength is defined as the likelihood of neurotransmitter release following an action potential (probability of release, P_r_). This probabilistic process is determined by the number of functional SV release sites and their individual probability of vesicle release. The probability of SV release is highly dependent on transient increases in intracellular Ca^2+^ levels at vesicular sensors. Accordingly, SV release sites and VGCCs are key substrates for generating diversity of synaptic function (Akbergenova et al. 2018; Aldahabi et al. 2022; Chen et al. 2015; Fedchyshyn and Wang 2005; Fekete et al. 2019; Gratz et al. 2019; Holderith et al. 2012; Laghaei et al. 2018; Miki et al. 2017; Nakamura et al. 2015; Newman et al. 2022; Rebola et al. 2019; Reddy-Alla et al. 2017; Sauvola et al. 2021; Sheng et al. 2012). Numerous studies have demonstrated that VGCC abundance is highly correlated with P_r_ across species (Akbergenova et al. 2018; Gratz et al. 2019; Holderith et al. 2012; Miki et al. 2017; Nakamura et al. 2015; Sheng et al. 2012). Paradoxically, this is not always the case. For example, a recent study investigated two cerebellar synaptic subtypes, one high-P_r_ formed by inhibitory stellate cells and one low-P_r_ formed by excitatory granule cells, and found higher VGCC levels at low-P_r_ granule synapses (Rebola et al. 2019). Since VGCCs in closer proximity to release sites are expected to have a greater impact on vesicular release probability than those positioned farther away, the spatial coupling of VGCCs and SVs at AZs is a critical determinant of P_r_ (Chen et al. 2015; Eggermann et al. 2011; Fedchyshyn and Wang 2005; Nakamura et al. 2015; Rebola et al. 2019). Indeed, at high-P_r_ stellate synapses, a ‘perimeter release’ AZ organization places VGCCs ∼40nm closer to SVs than at low-P_r_ granular synapses (Rebola et al. 2019). Another recent study investigated two functionally distinct connections formed by CA1 pyramidal cells (Aldahabi et al. 2022). While Ca^2+^ influx was higher at the high-P_r_ synapse, raising Ca^2+^ influx at the low-P_r_ synapse to match the high-P_r_ synapse did not equalize P_r_.

To further investigate this paradox, we sought a system where we could investigate the relationship between VGCCs and P_r_ both within and between two closely related neurons that form synapses with distinct release probabilities. *Drosophila* muscles are innervated by two glutamatergic motor neurons, one tonic and one phasic, that form type Ib and type Is synapses, respectively. Type Ib synapses have relatively low P_r_ and facilitate, whereas type Is synapses have higher P_r_ and depress in response to high-frequency stimulation (Aponte-Santiago et al. 2020; Lnenicka and Keshishian 2000). In this study, we investigated how VGCC abundance, spatial organization, and subunit composition contribute to synaptic heterogeneity at AZs of low-P_r_ type Ib and high-P_r_ type Is inputs to the same postsynaptic targets. We find that individual synapses formed by both low- and high-P_r_ inputs exhibit heterogeneous release properties that can be predicted by VGCC abundance alone. However, VGCC abundance does not correspond to differences in P_r_ between the two inputs. We identify underlying molecular and organizational differences that may alter the relationship between VGCC abundance and P_r_ at low- vs. high-P_r_ inputs. We further find that the homeostatic potentiation of neurotransmitter release triggered by glutamate receptor inhibition involves dynamic increases in VGCCs, the α2δ-3 subunit Straightjacket (Stj), and the AZ cytomatrix protein Bruchpilot (Brp) across AZs of both inputs. These findings provide insight into how VGCC and AZ protein abundance intersects with underlying molecular and organizational differences between inputs to contribute to greater synaptic diversity.

## RESULTS

### VGCC levels predict P_r_ within, but not between, inputs

To investigate the relationship between VGCC levels and neurotransmitter release properties at functionally distinct synapses, we took advantage of the two motor neuron subtypes with low and high release probabilities that innervate most *Drosophila* muscles (Aponte-Santiago and Littleton 2020; Kurdyak et al. 1994). These glutamatergic neuromuscular junctions (NMJs) contain hundreds of individual synapses that are accessible to single AZ functional imaging using genetically encoded Ca^2+^ indicators.

In *Drosophila*, Cacophony (Cac) is the sole Ca_v_2 pore-forming subunit and is the VGCC responsible for triggering synaptic transmission (Kawasaki, Felling, and Ordway 2000; Macleod et al. 2006; Peng and Wu 2007; Smith et al. 1996). To simultaneously monitor neurotransmitter release and VGCC levels, we swapped the N-terminal sfGFP tag in our well-characterized *cac^sfGFP-N^*line for a Td-Tomato tag (*cac^Td-Tomato-N^*) and confirmed that the tag does not impair synaptic function (Figs. 1A-F, S1; (Ghelani et al. 2023; Gratz et al. 2019)). We then expressed postsynaptically targeted GCaMP6f (SynapGCaMP6f; (Newman et al. 2017)), which reports Ca^2+^ influx through glutamate receptors in response to neurotransmitter release, in *cac^Td-Tomato-N^*animals for a plus/minus readout. We and others have previously shown that Cac levels are highly predictive of P_r_ at individual type Ib AZs (Akbergenova et al. 2018; Gratz et al. 2019). To determine if VGCC levels are similarly predictive at high-P_r_ type Is AZs, we measured Cac^Td-Tomato-N^ fluorescence intensity and monitored neurotransmitter release in response to 0.2-Hz stimulus at individual synapses. To enable direct comparisons between the two inputs, we simultaneously imaged type Ib and Is synapses at NMJ 6/7 (Fig. 1A). We quantified the number of times a vesicle was released over 120 stimuli to determine single-synapse P_r_. As has been previously reported, we found that type Is synapses exhibited significant heterogeneity and higher average P_r_ than type Ib synapses (Fig. 1B, C; (Lu et al. 2016; Newman et al. 2022)). Consistent with their higher P_r_, type Is connections contain relatively fewer low-P_r_ and more high-P_r_ AZs (Figs. 1D). We next investigated the correlation between P_r_ and VGCC levels and found that at type Is inputs, single-AZ Cac intensity positively correlates with P_r_ (Fig. 1E). We also observe a strong positive correlation between VGCC levels and P_r_ at type Ib inputs to the same muscles (Fig. 1F), consistent with our and others’ prior findings (Akbergenova et al. 2018; Gratz et al. 2019; Newman et al. 2022).

**Figure 1:**
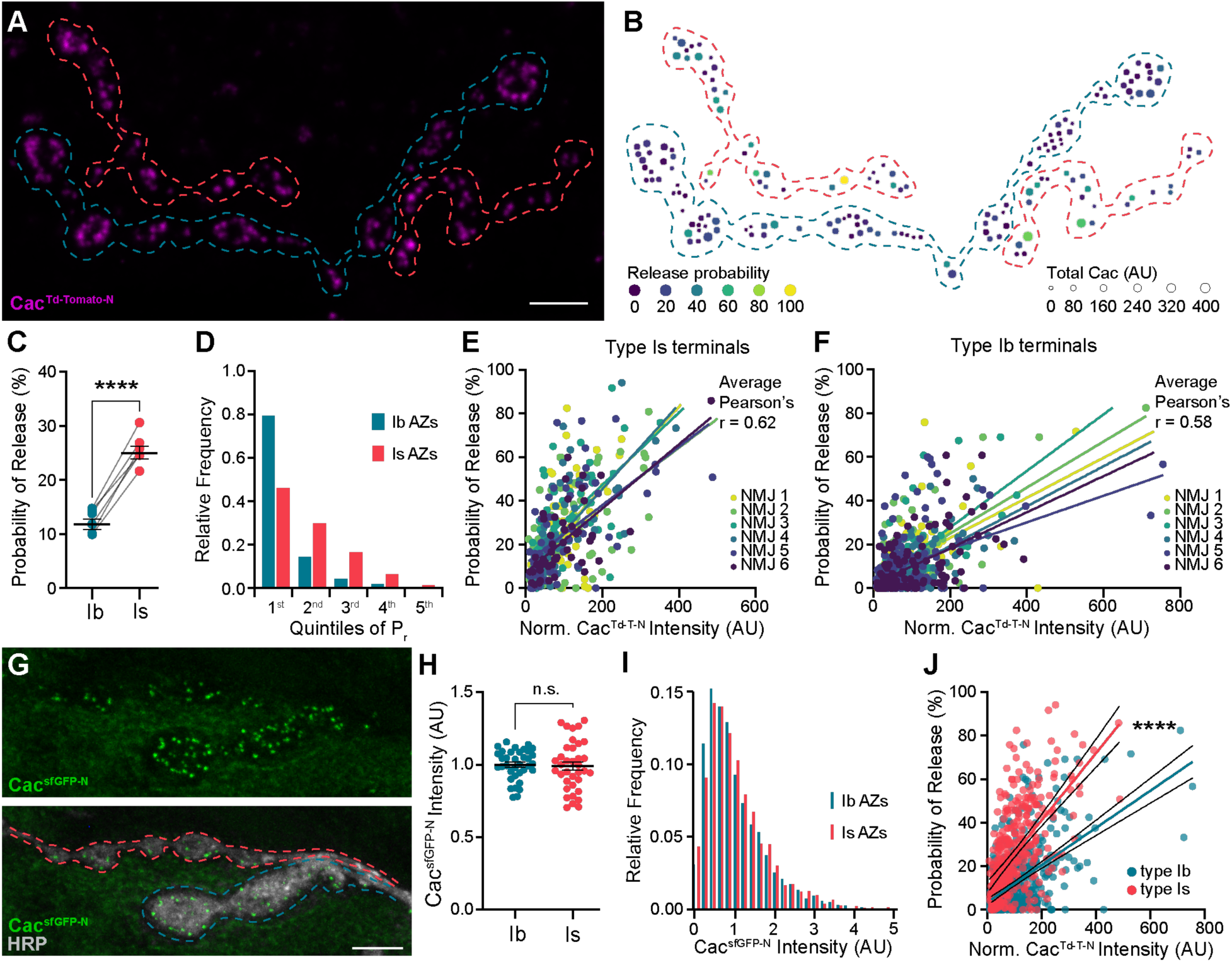
VGCC levels predict Pr within, but not between, inputs. **(A)** Representative confocal Z-projection of Cac^Td-Tomato-N^ (magenta) with type Ib (blue) and type Is (red) terminals outlined. **(B)** AZ heat map of terminals in A with color indicating Pr and size representing sum Cac intensity levels in arbitrary units (AU). **(C)** Average single-AZ probability of release at type Ib and Is terminals. **(D)** Quintile distribution of single-AZ Pr frequency at type Ib and Is inputs. **(E, F)** Correlation between normalized Cac^Td-Tomato-N^ intensity and Pr at type Is and Ib AZs of the same 6 NMJs. Each dot represents a single AZ and each color corresponds to an individual NMJ with linear regression lines indicated for each. **(G)** Top, representative confocal Z-projection of Cac^sfGFP-N^. Bottom, Cac^sfGFP-N^ in green with HRP marking neuronal membranes in gray. Type Ib (blue) and type Is (red) terminals are outlined. **(H)** Quantification of Cac^sfGFP-N^ AZ intensity at type Ib and Is terminals. Each data point represents the average normalized single AZ sum intensity for an individual NMJ. **(I)** Distribution of normalized Cac^sfGFP-N^ intensity from single type Ib and Is AZs in H (X-axis cutoff at 5.0). **(J)** Comparison between normalized Cac^Td-Tomato-N^ and Pr of type Ib and Is AZs combined from E-F with linear regression lines (blue and red, respectively) and 95% confidence intervals (black lines) indicated. All scale bars = 5µm.

A simple prediction of the observation that VGCC levels correlate highly with P_r_ at individual AZs of both low- and high-P_r_ inputs is that Cac levels will be higher at synapses of type Is inputs than type Ib. We analyzed Cac^sfGFP-N^ levels at individual type Ib and Is synapses and found that average Cac levels are the same at type Ib and Is AZs (Fig. 1G, H). Cac levels are also similarly distributed across AZs of the two inputs (Fig. 1I). Together, these findings indicate that the relationship between VGCC levels and P_r_ differs between the two inputs. Consistently, when we directly compare the best-fit lines for the relationship between Cac levels and P_r_ at type Ib and Is inputs from our correlative functional imaging data (Fig. 1E, F), we find that the slopes are significantly different (Fig. 1J). Across type Is AZs, a similar range of VGCC levels supports a higher range of release probabilities. Thus, VGCCs can predict P_r_ within synaptic subtypes, but not between AZs of different synaptic subtypes, providing a framework for understanding seemingly contradictory findings on the role of VGCCs in determining P_r_.

### VGCC clusters are more compact at AZs of high-P_r_ type Is inputs

Many differences between low-P_r_ type Ib and high-P_r_ type Is AZs have been described (Aponte-Santiago and Littleton 2020; Aponte-Santiago et al. 2020; Atwood, Govind, and Wu 1993; He et al. 2023; Jetti et al. 2023; Kurdyak et al. 1994; Lu et al. 2016; Medeiros and O’Connor-Giles 2023).

Perhaps most notably, type Is AZs experience ∼2-fold greater Ca^2+^ influx than type Ib (He et al. 2023; Lu et al. 2016). While this alone could explain the estimated 3-fold greater P_r_ at type Is AZs and is certainly a key factor, several lines of evidence argue for additional contributors. A recent study using a botulinum transgene to isolate type Ib and Is synapses for electrophysiological analysis found that increasing external [Ca^2+^] from physiological levels (1.8 mM) to 3 mM or even 6 mM does not result in a 3-fold increase in EPSCs or quantal content at type Ib synapses and type Ib synapses continue to facilitate at 3 mM external [Ca^2+^] (He et al. 2023). Using this approach, they further found that type Ib synapses are more sensitive to the slow Ca^2+^ chelator EGTA, indicating looser VGCC-SV coupling.

We investigated the spatial distribution of VGCCs at type Ib and Is AZs using 3D dSTORM single-molecule localization microscopy (SMLM). An individual VGCC complex is estimated to be ∼10 nm in diameter with the most common immunolabeling techniques adding significantly to their size and creating a linkage error of ∼20 nm between the target molecule and fluorescent reporter (Früh et al. 2021; Liu, Hoess, and Ries 2022; Thomas 2000). For following VGCC dynamics using single-particle tracking via photoactivation localization microscopy (sptPALM), we recently incorporated mEOS4b (Paez-Segala et al. 2015) at the same N-terminal site we previously used to endogenously tag Cac, achieving a linkage error of less of than 5 nm (Ghelani et al. 2023; Gratz et al. 2019). To gain more flexibility in labeling Cac without adding to the linkage error, we swapped the mEOS tag for a similarly sized HaloTag (*cac^HaloTag-N^*). *cac^HaloTag-N^* flies are fully viable, do not display significant defects in synaptic function, and exhibit normal Cac localization at AZs as observed in super-resolution optical reassignment images, where Brp is arranged in rings surrounding puncta of VGCCs (Fig. S1; 2A-C, (Ghelani et al. 2023)). HaloTag, which covalently binds synthetic ligands, is 3.3 nm in diameter (Los et al. 2008; Yazaki et al. 2019), yielding a linkage error well under 5 nm.

*cac^HaloTag-N^* larvae were stained with JaneliaFluor646 HaloTag ligand (Grimm et al. 2015) and horseradish peroxidase (HRP) to distinguish between type Ib and Is branches and enable simultaneous imaging of the two inputs at a single NMJ. We then used density-based spatial clustering of applications with noise (DBSCAN) analysis to identify Cac clusters at type Ib and Is AZs (Fig. 2D,E; (Ehmann et al. 2014)). We find that the average size of Cac^HaloTag-N^ clusters is similar at low- and high-P_r_ AZs (Fig. 2F), with mean diameters of approximately 102 nm and 105 nm, respectively. This is similar to the Cac^mEOS4b-N^ type Ib cluster size observed by sptPALM imaging (Ghelani et al. 2023). In agreement with our confocal level data, the number of localizations per cluster was similar at low- and high-P_r_ AZs (Fig. 2G). We then calculated the average Cac density per AZ and found that VGCCs are significantly more densely organized at high-P_r_ type Is AZs than low-P_r_ type Ib AZs (Fig. 2H, I). Greater VGCC AZ density at type Is AZs is consistent with a recent SMLM study using antibodies to label Cac, indicating these results are robust to different labeling approaches (Newman et al. 2022). Together, these findings suggest that more compact organization of VGCCs increases their coupling to SVs and contributes to the steeper relationship between VGCC levels and P_r_ at high-P_r_ type Is AZs.

**Figure 2:**
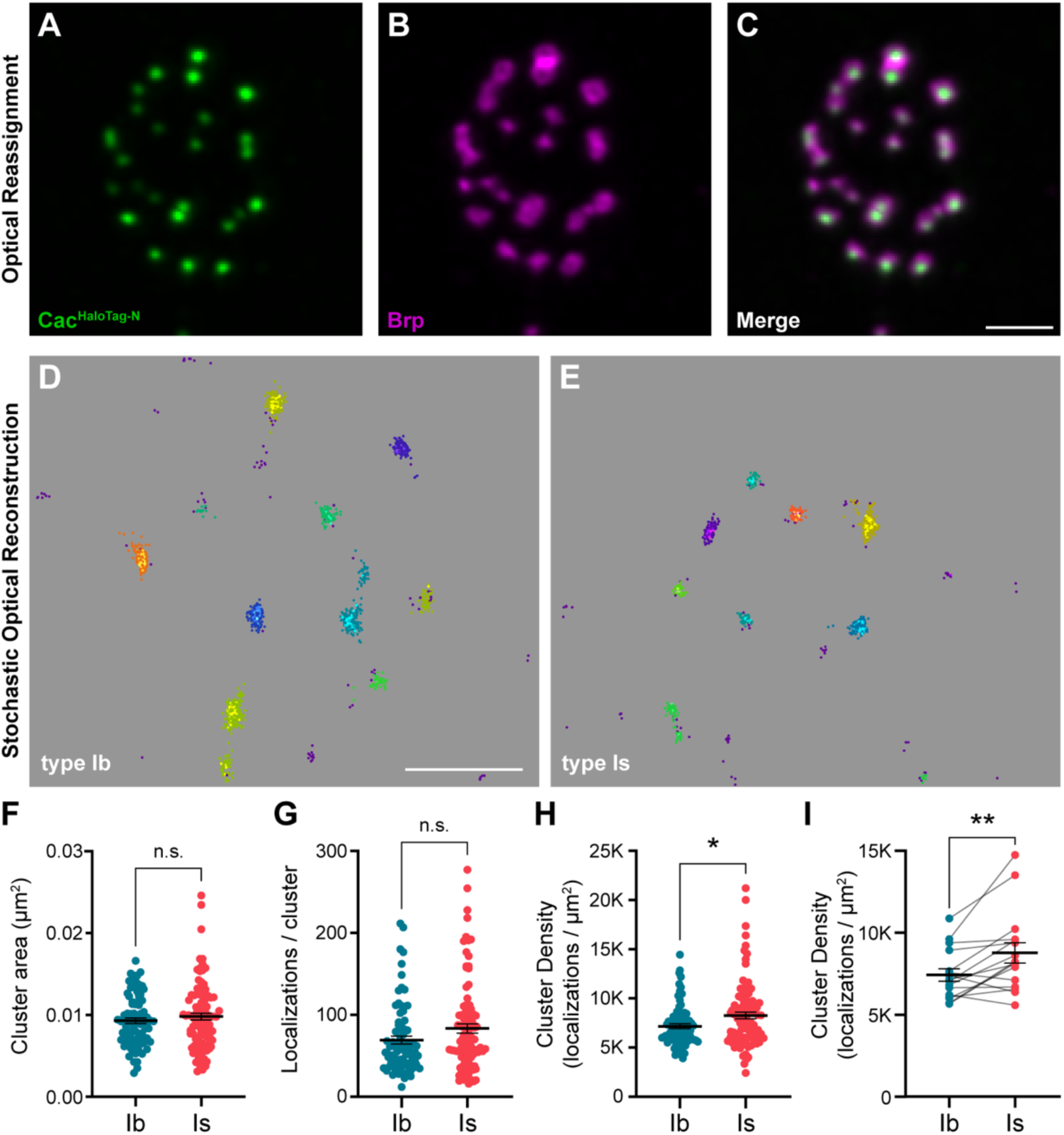
VGCC clusters are more compact at AZs of high-Pr type Is inputs. **(A-C)** Representative SoRa Z-projection of Cac^HaloTag-N^ (green), Brp (magenta), and merge. **(D, E)** Representative boutons of STORM Cac^HaloTag-N^ clusters as identified by DBSCAN at type Ib and Is boutons as indicated. Each color represents an individual identified cluster with purple scattered dots identifying excluded background signal. **(F-H)** Analysis of STORM-acquired Cac^HaloTag-N^ clusters where each data point represents the respective single-cluster measurement averaged over individual boutons. **(F)** Quantification of Cac^HaloTag-N^ cluster area at type Ib and Is AZs. **(G)** Quantification of localizations per cluster at type Ib and Is boutons. **(H)** Calculated Cac^HaloTag-N^ cluster density at type Ib and Is AZs. **(I)** Paired analysis of calculated AZ cluster density averaged over individual type Ib and Is inputs to the same muscle. All scale bars = 1µm.

### Differences in Bruchpilot levels and function at low- and high-P_r_ inputs

To understand how these nanoscale differences in VGCC organization might be established, we investigated the AZ scaffolding protein Brp. Brp/CAST/ELKS family proteins function as central organizers of both VGCCs and SV release sites at developing synapses (Dai et al. 2006; Dong et al. 2018; Hallermann et al. 2010; Held, Liu, and Kaeser 2016; Kittel et al. 2006; Liu et al. 2014; McDonald, Fetter, and Shen 2020; Radulovic et al. 2020). Like Cac, Brp is more densely arranged at type Is AZs as measured through SMLM and stimulated emission depletion (STED) imaging studies (He et al. 2023; Jetti et al. 2023; Mrestani et al. 2021). We simultaneously imaged type Ib and Is inputs and found lower Brp levels at type Is AZs (Fig. 3A, B). Since Cac levels are similar at AZs of the two inputs, lower Brp levels result in a significantly higher Cac:Brp ratio at type Is synapses, which we hypothesize promotes compact organization of VGCCs (Fig. 3C). In contrast, we and others have previously shown that Brp levels positively correlate with P_r_ among AZs of low-P_r_ type Ib inputs (Gratz et al. 2019; Muhammad et al. 2015; Newman et al. 2017; Peled, Newman, and Isacoff 2014; Reddy-Alla et al. 2017). Consistently, Brp and Cac levels strongly correlate at type Ib AZs (Gratz et al. 2019) and we observe a similarly strong correlation across individual type Is AZs (Fig. 3D). Thus, like VGCCs, Brp levels contribute in distinct ways to synaptic heterogeneity within vs. between low- and high-P_r_ inputs, likely due to differences in AZ organization between the two synaptic subtypes.

**Figure 3:**
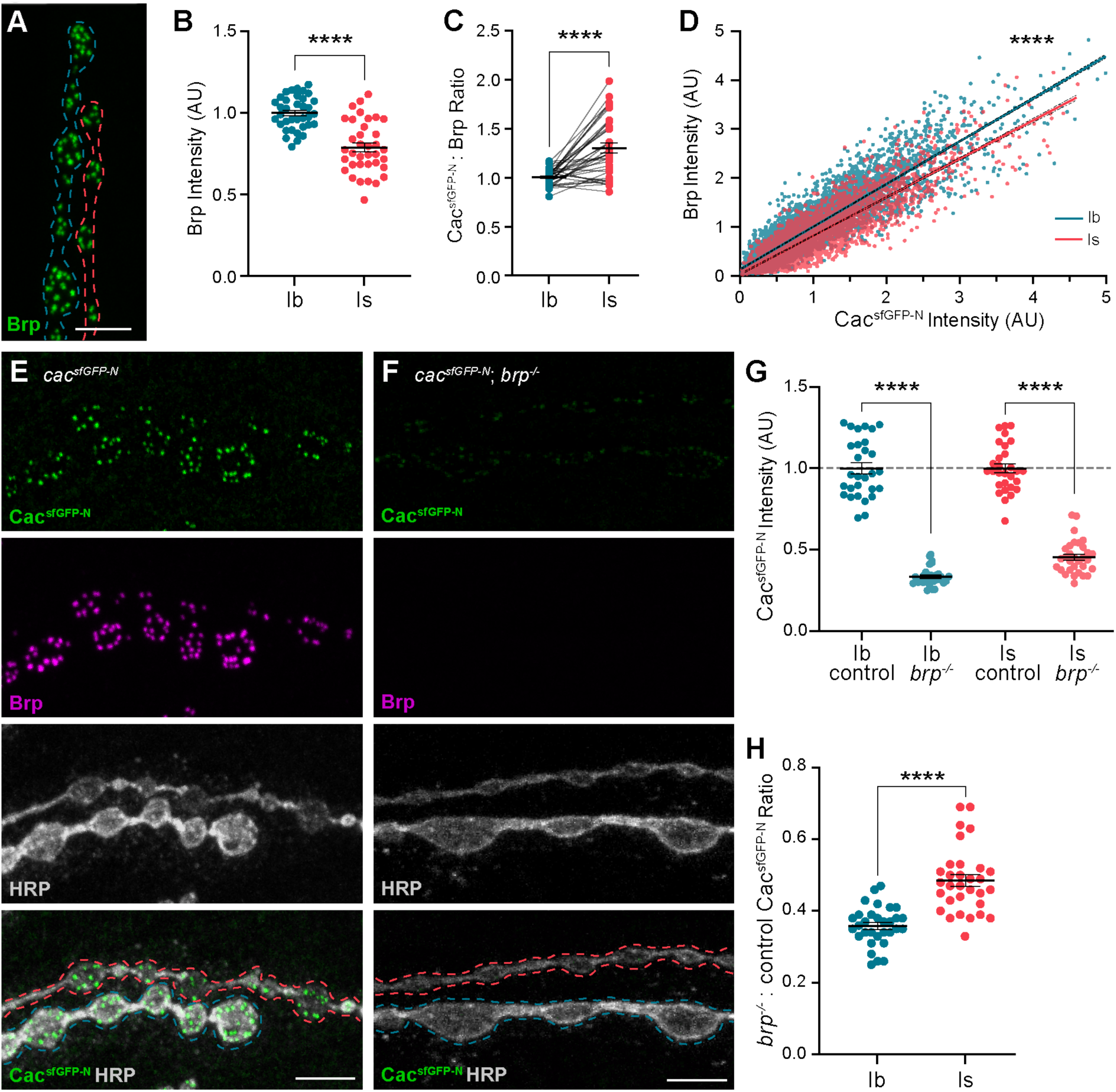
Differences in Bruchpilot (Brp) levels and function at low-and high-Pr inputs. **(A)** Representative confocal Z-projection of Brp expression at type Ib (blue outline) and type Is (red outline) terminals. **(B)** Quantification of Brp AZ intensity at type Ib and Is terminals. **(C)** Ratio of normalized Cac^sfGFP-N^:Brp levels at type Ib and Is inputs to the same muscles. **(D)** Correlation of Cac^sfGFP-N^ and Brp at type Ib and Is single AZs with linear regression lines (blue and red, respectively) and 95% confidence intervals (black dotted lines) indicated. **(E, F)** Representative confocal Z-projections of Cac^sfGFP-N^ (green), Brp (magenta), HRP (white), and merge at type Ib (blue outline) and Is (red outline) terminals of *cac^sfGFP-N^* (control) or *cac^sfGFP-N^*;*brp^-/-^* (*brp^-/-^*) animals. **(G)** Quantification of Cac^sfGFP-N^ normalized fluorescence intensity at type Ib and Is AZs of control vs *brp^-/-^* NMJs. **(H)** Ratio of Cac^sfGFP-N^ fluorescence intensity at type Ib and Is AZs between *brp^-/-^*and control NMJs. For B and G, each data point represents the average normalized single AZ sum intensity for an individual NMJ. All scale bars = 5µm.

We next investigated the requirement for Brp in promoting VGCC accumulation at low- and high-P_r_ inputs by analyzing Cac^sfGFP-N^ levels in *brp* null mutants (*brp^-/-^*; Fig. 3E, F). Cac^sfGFP-N^ levels are diminished at both type Ib and Is AZs, demonstrating a conserved role for Brp in promoting Cac accumulation at both inputs (Fig. 3G). The relative decrease in Cac levels at type Ib AZs is significantly greater than at type Is AZs, indicating a greater requirement for Brp in regulating VGCC levels at low-P_r_ type Ib synapses (Fig. 3H). This suggests that an additional factor or factors function with or upstream of Brp to establish differences in Brp dependence at low- and high-P_r_ AZs.

### Brp differentially regulates VGCC dynamics at low- and high-P_r_ synapses during presynaptic homeostatic potentiation

In response to acute or chronic inhibition of glutamate receptors at NMJs, *Drosophila* motor neurons homeostatically increase neurotransmitter release to maintain synaptic communication (Davis and Muller 2015; Frank 2014; James, Zwiefelhofer, and Frank 2019). Pharmacological inhibition of glutamate receptors with the wasp toxin Philanthotoxin-433 (PhTx) induces acute presynaptic homeostatic potentiation of release (PHP) within minutes (Frank et al. 2006). We and others have demonstrated that acute PHP involves rapid changes in VGCCs and other AZ protein levels at type Ib AZs (Bohme et al. 2019; Gratz et al. 2019; Weyhersmuller et al. 2011). Recent studies have revealed significant differences in the induction of PHP at low- and high-P_r_ synaptic inputs under different conditions (Genc and Davis 2019; Newman et al. 2017; Sauvola et al. 2021). PhTx induces acute PHP at both type Ib and Is synapses (Genc and Davis 2019), but the molecular changes underlying PHP at high-P_r_ type Is AZs remain unknown. To compare the dynamic modulation of VGCCs at low- and high- P_r_ AZs, we treated *cac^sfGFP-N^* larvae with non-saturating concentrations of PhTx for 10 minutes, then quantified Cac and Brp levels at type Ib and Is AZs (Fig. 4A, B). We observe a significant PhTx-induced increase in Brp and Cac^sfGFP-N^ levels at type Is AZs similar to type Ib (Fig. 4C, D; (Gratz et al. 2019)). Thus, despite their distinct baseline transmission and organizational properties, PhTx-induced potentiation of neurotransmitter release involves the rapid accumulation of VGCCs at both low- and high-P_r_ AZs.

**Figure 4:**
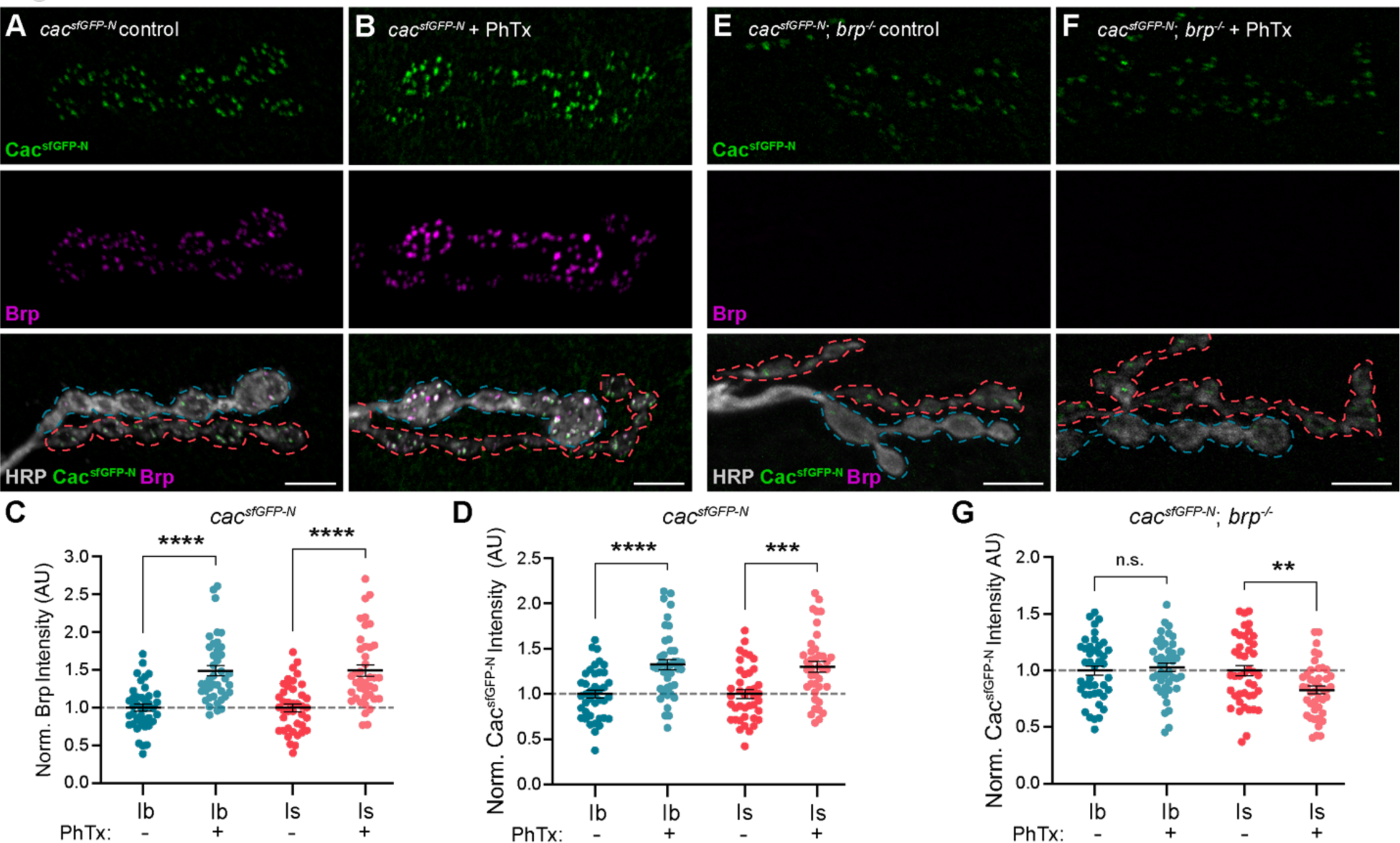
Brp differentially regulates VGCC dynamics at low-and high-Pr inputs during presynaptic homeostatic potentiation. **(A, B)** Representative confocal Z-projections of Cac^sfGFP-N^ (top, green), Brp (middle, magenta), and both merged with HRP (bottom, gray) at untreated and PhTx-treated *cac^sfGFP-N^* NMJs showing type Ib (blue) and type Is (red) terminals. **(C)** Quantification of Brp fluorescence intensity at untreated and PhTx-treated type Ib and Is terminals. **(D)** Quantification of Cac^sfGFP-N^ fluorescence intensity at untreated and PhTx-treated type Ib and Is terminals. **(E, F)** Representative confocal Z-projections of Cac^sfGFP-N^ (top, green), Brp (middle, magenta), and both merged with HRP (bottom, gray) at untreated and PhTx-treated *cac^sfGFP-N^*;*brp^-/-^*NMJs showing type Ib (blue) and type Is (red) terminals. **(G)** Quantification of Cac^sfGFP-N^ fluorescence intensity at untreated and PhTx-treated *cac^sfGFP-N^*;*brp^-/-^* type Ib and Is terminals. For all quantifications, each data point represents the average normalized single AZ sum intensity for an individual NMJ. All scale bars = 5µm.

At low-P_r_ type Ib AZs, Brp is a critical regulator of PHP-induced accumulation of proteins associated with SV priming and release, specifically Unc13A and Syntaxin-1A (Bohme et al. 2019). At type Ib AZs, PhTx also induces a Brp-dependent increase in Cac density and decrease in channel mobility (Ghelani et al. 2023). Notably, Brp itself becomes more densely organized during PHP (Ghelani et al. 2023), consistent with its denser organization at high-P_r_ type Is AZs (Mrestani et al. 2021). Since baseline accumulation of VGCCs depends less on Brp at high-P_r_ type Is AZs, we investigated the role of Brp in promoting dynamic increases in VGCC levels at type Ib and Is AZs by treating *cac^sfGFP-N^; brp^-/-^* larvae with PhTx followed by quantification of Cac^sfGFP-N^ levels (Fig. 4E, F). We find that PhTx failed to induce accumulation of Cac at either type Ib or Is AZs in *brp^-/-^* mutants, demonstrating a shared requirement for Brp in regulating VGCC dynamics at both inputs (Fig. 4G). In contrast to no change at type Ib AZs, Cac^sfGFP-N^ levels are significantly decreased at type Is AZs (Fig. 4G), revealing an input-specific role for Brp in maintaining VGCC levels during the dynamic reorganization of AZs. Consistently, Ghelani et al. (2023) found that whereas PhTx induces a decrease in the Cac mobility at wild-type type Ib AZs, in *brp^-/-^* mutants Cac mobility increases (Ghelani et al. 2023). Together, these findings suggest potentiating synapses must coordinate the accumulation of new VGCCs with the stabilization of existing channels, and that meeting this challenge is more dependent upon Brp at high-P_r_ AZs.

### Endogenous tagging of VGCC auxiliary subunits reveals distinct synaptic expression patterns

In addition to the pore-forming α subunits, VGCCs comprise auxiliary α2δ and β subunits that regulate forward channel trafficking, membrane insertion, and function (Fig. 5A; (Campiglio and Flucher 2015; Dolphin and Lee 2020; Weiss and Zamponi 2017)). β subunits interact with pore-forming α subunits intracellularly, whereas GPI-anchored α2δ subunits are largely extracellular. Beyond their interaction with α subunits, α2δs have been shown to interact with a growing number of extracellular proteins to promote synaptogenesis (Bauer et al. 2010; Dolphin 2018; Risher et al. 2018). The *Drosophila* genome encodes one synaptic Ca_v_2 α subunit (Cac), one β subunit, and three α2δ subunits (Littleton and Ganetzky 2000). Auxiliary subunits are both spatially and temporally regulated and broadly able to interact with α subunits. Thus, the subunit composition of channel complexes is a potential source of significant diversity in both the spatial and functional regulation of VGCCs.

**Figure 5:**
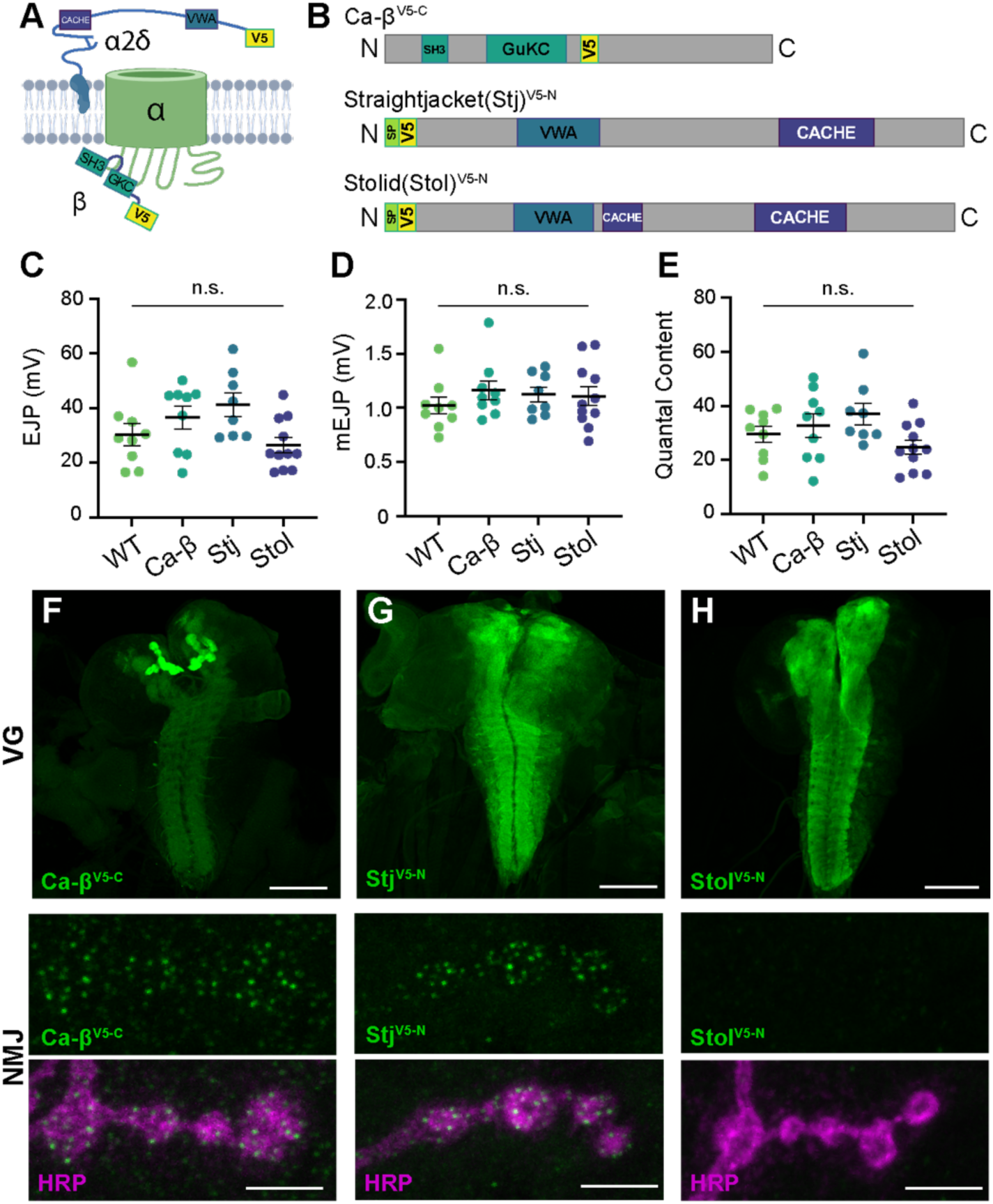
Endogenous tagging of VGCC auxiliary subunits reveals distinct synaptic expression patterns. **(A)** Schematic of a Ca^2+^ channel complex with tagged auxiliary subunits (created with BioRender). **(B)** Schematic of Ca-β (isoform PL shown), Stj (isoform PC), and Stolid (isoform H/I) indicating endogenous tag locations. **(C-E)** Quantification of EJPs, mEJPs, and quantal content for each endogenously tagged line. **(F-H)** Representative confocal Z-projections of auxiliary subunit expression (green) at the larval ventral ganglion (VG, top, scale bars = 100µm) and NMJs co-labeled with anti-HRP (magenta, bottom, scale bars = 5µm).

*Ca-β* encodes the sole *Drosophila* β subunit and has been shown to enhance Ca^2+^ transients in sensory neurons (Kanamori et al. 2013). *Drosophila* α2δ-3, also known as Straightjacket (Stj), has well-characterized roles at the NMJ in promoting Ca^2+^ channel clustering, homeostatic plasticity, and, independently of Cac, synapse formation and organization (Dickman, Kurshan, and Schwarz 2008; Hoover et al. 2019; Kurshan, Oztan, and Schwarz 2009; Ly et al. 2008; Schöpf et al. 2021; Wang et al. 2016). While low sequence homology between α2δ subunits within and across species makes 1:1 mapping difficult, the remaining two *Drosophila* α2δs map more closely to mammalian α2δ-3 and -4 than α2δ-1 and -2. Stolid was recently shown to promote dendritic Cac expression in motor neurons, whereas Ma2d is known to function in muscle where it is broadly expressed (Heinrich and Ryglewski 2020; Reuveny et al. 2018). The synaptic localization of endogenous auxiliary subunits with VGCCs remains unknown in *Drosophila*.

To explore potential differences in VGCC subunit composition at type Ib and Is synapses, we used CRISPR gene editing to incorporate endogenous V5 tags in sequence common to all isoforms of *stj*, *stolid,* and *Ca-β* (Bruckner et al. 2017; Gratz et al. 2014). We inserted V5 after the N-terminal signal peptides of Stj and Stolid and near the C-terminus of Ca-β (see Materials and Methods for details), and confirmed that the incorporation of the peptide tags did not impair neurotransmission (Fig. 5B-E). We investigated the expression of each endogenously tagged subunit in the larval ventral ganglion and found that all subunits are expressed in the synaptic neuropil in a pattern similar to the α subunit Cac (Fig. 5F-H; (Gratz et al. 2019)). Similar to Cac, Ca-β^V5-C^ is highly enriched in the mushroom bodies of the larval brain. We next investigated expression at the larval NMJ where Cac localizes in a single punctum at each AZ and found that only Ca-β^V5-C^ and Stj^V5-N^ are present (Fig. 5F-H). This aligns with the recent finding that Stolid does not play a role in regulating Ca^2+^ transients at the larval NMJ (Heinrich and Ryglewski 2020). We also observe Ca-β^V5-C^ expression in muscle as expected for the sole *Drosophila* β subunit (Fig. 5F and see Fig. 6C).

**Figure 6:**
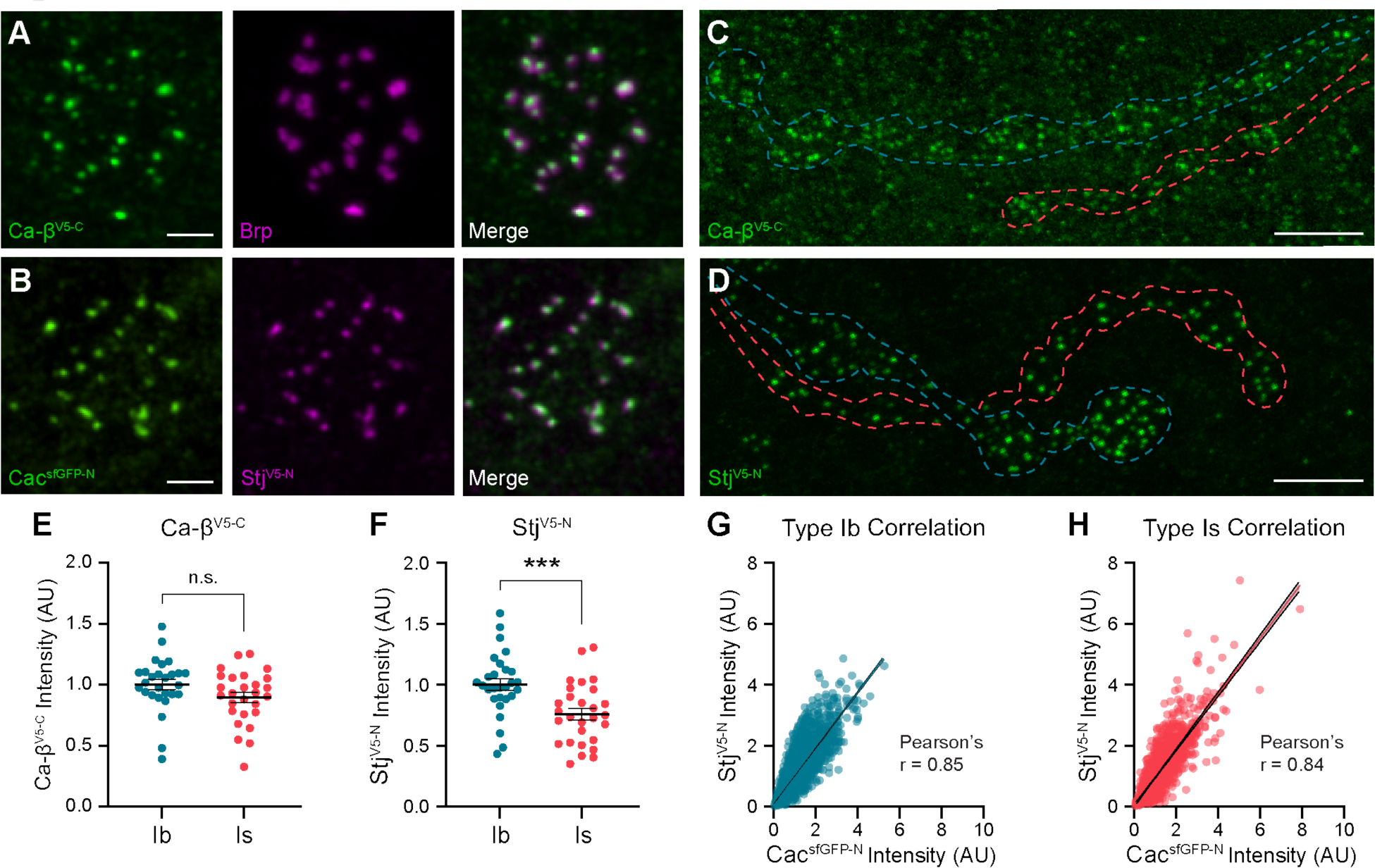
Stj/α2δ-3 levels are lower at AZs of high-Pr type Is inputs. **(A)** Representative SoRa Z-projections of Ca-β^V5-C^ (green), Brp (magenta), and merge at a single bouton. **(B)** Representative SoRa Z-projections of Cac^sfGFP-N^ (green), Stj^V5-N^ (magenta), and merge at a single bouton. Scale bars for A and B = 1μm. **(C, D)** Representative confocal Z-projections of Ca-β^V5-C^ expression and Stj^V5-N^ expression at type Ib (blue outline) and type Is (red outline) terminals. Scale bars = 5μm. **(E, F)** Quantification of Ca-β^V5-C^ and Stj^V5-N^ fluorescence intensity at type Ib and Is AZs. Each data point represents the average normalized single AZ sum intensity for an individual NMJ. **(G, H)** Correlation of Cac^sfGFP-N^ and Stj^V5-N^ fluorescence intensity levels at type Ib and Is single AZs with linear regression lines (blue or red line, respectively) and 95% confidence intervals (black lines).

### Stj/α2δ-3 levels are lower at AZs of high-P_r_ type Is inputs

To investigate Ca-β^V5-C^ and Stj^V5-N^ localization at type Ib and Is AZs, we used super-resolution optical reassignment microscopy. Both subunits localize to AZs labeled with Cac or the CAST/ELKS AZ cytomatrix protein Brp (Fig. 6A, B). We observe Brp rings surrounding puncta of VGCCs including Ca-β^V5-C^ (Fig. 6A). The tight localization of both subunits to central AZ puncta suggests they are associated with α subunits and predicts that Ca-β^V5-C^ and Stj^V5-N^ levels, like Cac, will be similar at the low- and high-P_r_ synapses. To test this, we simultaneously imaged Ca-β^V5-C^ and Stj^V5-N^ levels at both inputs using confocal microscopy and measured fluorescence intensity (Fig. 6C, D). As predicted, we found that Ca-β^V5-C^ levels are similar at type Ib and Is AZs (Fig. 6E). In contrast, Stj^V5-N^ levels are significantly lower at high-P_r_ type Is AZs (Fig. 6F). Thus, while Cac and Ca-β are present in similar ratios at AZs of both inputs, surprisingly, the same is not true of Stj/α2δ-3 with high-P_r_ type Is AZs exhibiting lower levels of Stj. This unexpected finding indicates that α:α2δ-3 stoichiometry is not always 1:1 *in vivo* and differs at low- and high-P_r_ synapses. This is consistent with studies of mammalian subunits indicating that in contrast to β subunits, α2δ interactions with α subunits may be transient, leading to a pool of VGCCs lacking α2δ (Muller et al. 2010; Voigt et al. 2016). Our results indicate this pool may be present *in vivo* and larger at high-P_r_ type Is inputs.

To further investigate the contribution of Stj to synaptic heterogeneity, we analyzed the relationship between Cac and Stj levels at individual AZs of type Is inputs. Stj^V5-N^ and Cac^sfGFP-N^ levels are highly positively correlated at type Is AZs (Fig. 6G). We observe the same relationship between Stj^V5-N^ and Cac^sfGFP-N^ levels at type Ib AZs (Fig. 6H). Because P_r_ is highly positively correlated with Cac levels within synaptic subtypes, this indicates that Stj levels are also positively correlated with P_r_ within, but not between, inputs.

### Stj/α2δ-3 levels are modulated at AZs of both low- and high-P_r_ inputs during presynaptic homeostatic potentiation

α2δ subunits are critical regulators of α subunit forward trafficking. In flies and mammals, overexpression of α2δ subunits increases α subunit abundance, whereas overexpression of the α subunit alone does not (Cao et al. 2004; Cunningham et al. 2022; Hoppa et al. 2012). These findings suggest that α2δ may be dynamically regulated together with Cac during PHP, a prediction we can now test with our endogenously tagged line. Following PhTx exposure, we find that Stj^V5-N^ is recruited on a rapid timescale to both low- and high-P_r_ AZs, increasing by a similar percentage at both type Ib and Is AZs (27% and 26%, respectively) as predicted (Fig. 7A-C). Cac levels are similarly increased (33% at type Ib and 30% at type Is; See Fig. 4D), suggesting coordinated regulation. We have previously shown that Cac abundance is also increased in chronic PHP, which is induced by genetic loss of the GluRIIA receptor subunit (Gratz et al. 2019; Li et al. 2018; Petersen et al. 1997). We investigated Stj dynamics in *GluRIIA^-/-^* null animals and observed elevated Stj^V5-N^ levels at AZs of both type Ib and Is inputs (18% and 17%, respectively; Fig. 7D-F), indicating similar dynamics during chronic PHP. These data are consistent with the recent finding that in addition to its previously uncovered role in promoting an increase in the readily releasable pool of SVs (Wang et al. 2016), Stj is required for the accumulation of Cac at type Ib AZs during both acute and chronic PHP (Zhang et al. 2023). Thus, the abundance of multiple key AZ proteins distinguishes low- and high-P_r_ synapses within, but not between, inputs at baseline and during homeostatic plasticity.

**Figure 7:**
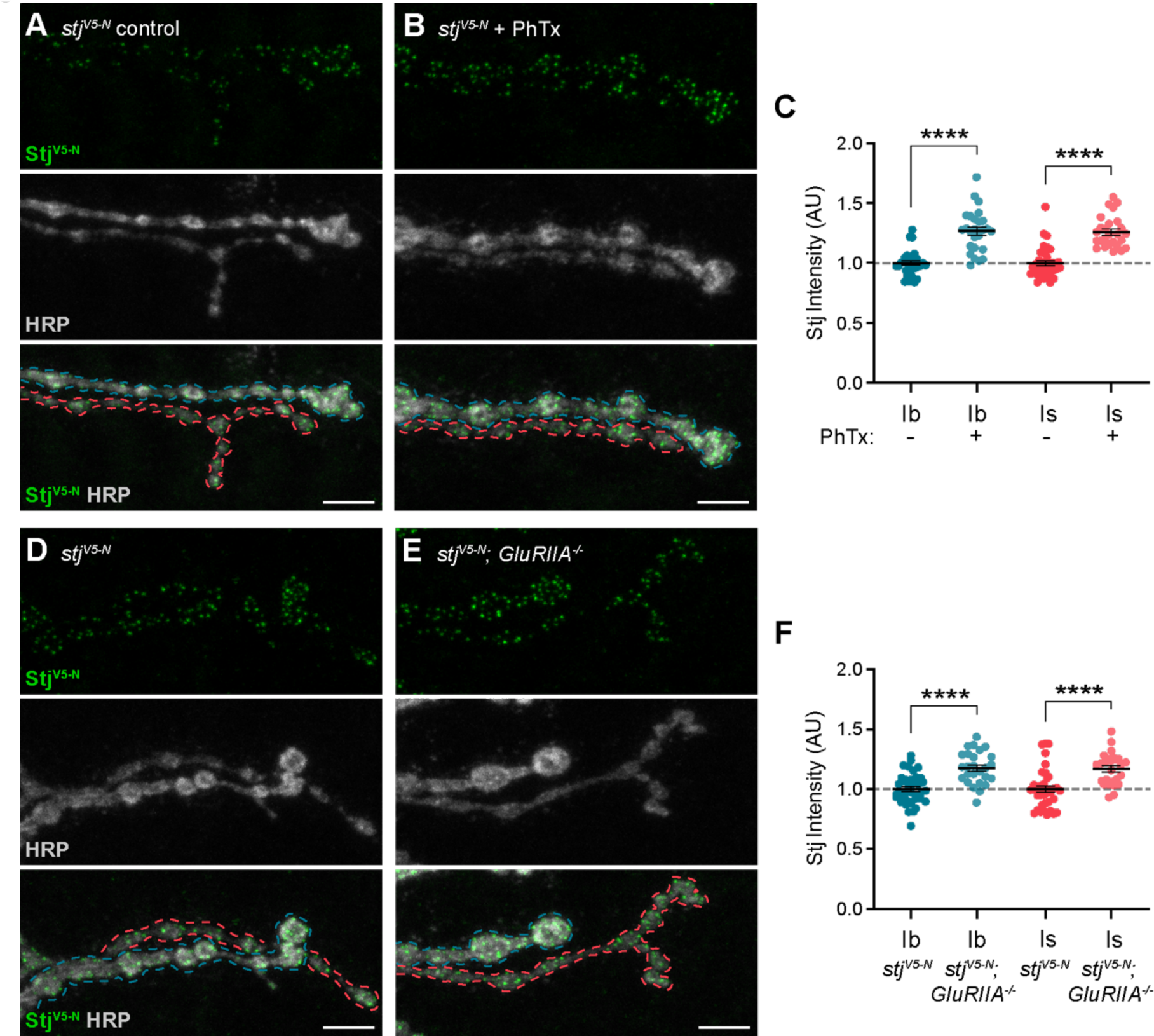
Stj/α2δ-3 levels are modulated at AZs of both low- and high-Pr inputs during presynaptic homeostatic potentiation. **(A, B)** Representative confocal Z-projections of Stj^V5-N^ (top, green), HRP (middle, gray), and merge (bottom) at untreated and PhTx-treated *stj^V5-N^* NMJs showing type Ib (blue) and type Is (red) terminals. **(C)** Quantification of Stj^V5-N^ fluorescence intensity at untreated and PhTx-treated type Ib and Is terminals. **(D-E)** Representative confocal Z-projections of Stj^V5-N^ (top, green), HRP (middle, gray), and merge (bottom) at *stj^V5-N^* and *stj^V5-N^*;*GluRIIA^-/-^*NMJs showing type Ib (blue) and type Is (red) terminals. **(F)** Quantification of Stj^V5-N^ fluorescence intensity levels at *stj^V5-N^* and *stj^V5-N^*;*GluRIIA^-/-^* type Ib and Is terminals. For all quantifications, each data point represents the average normalized single AZ sum intensity for an individual NMJ. All scale bars = 5µm.

## DISCUSSION

Complex nervous system function depends on communication at synapses with heterogeneous and plastic properties. Paradoxical findings in the field have raised questions about the role of VGCCs in establishing neurotransmitter release properties. Our findings suggest a model in which two broad intersecting mechanisms contribute to synaptic diversity in the nervous system: (1) nanoscale spatial organization and relative molecular content establish distinct average basal release probabilities that differ between inputs and (2) coordinated modulation of VGCC and active zone protein abundance independently tunes P_r_ among individual synapses of distinct inputs. This model provides a framework for integrating diverse findings in the field and understanding how multiple levels of molecular and organizational diversity can intersect to generate extensive synaptic heterogeneity.

Investigations at diverse synapses using approaches ranging from cell-attached patch recordings to freeze-fracture immuno-electron microscopy to correlative functional imaging have revealed a strong positive correlation between VGCC number and P_r_ (Akbergenova et al. 2018; Gratz et al. 2019; Holderith et al. 2012; Miki et al. 2017; Nakamura et al. 2015; Newman et al. 2022; Sheng et al. 2012). This holds true among mature synapses in the hippocampus or immature synapses of the calyx of Held (Holderith et al. 2012; Sheng et al. 2012) and at the developing *Drosophila* NMJ where the differences in P_r_ and channel number between AZs of a single input also correlate with synapse maturity (Akbergenova et al. 2018; Gratz et al. 2019; Newman et al. 2022). While this conclusion corresponds neatly with the dependence of neurotransmitter release on Ca^2+^ influx, counterintuitively, a disconnect between VGCC and P_r_ is observed in some studies. Although the number of functional VGCCs positively correlates with P_r_ among synapses in the immature calyx of Held, the mature calyx has higher P_r_ yet recruits fewer Ca^2+^ channels (Fedchyshyn and Wang 2005; Sheng et al. 2012; Wang and Augustine 2014). Similarly, in the cerebellum, inhibitory stellate cells form high-P_r_ synapses with lower VGCC levels than low-P_r_ synapses formed by excitatory granular cells (Rebola et al. 2019). Adding to these paradoxical examples, we find that VGCC levels positively correlate with P_r_ among the heterogeneous synapses formed by either low-P_r_ type Ib or high-P_r_ type Is motor neurons, but overall VGCC abundance is similar at the two inputs (Figs. 1, 2) despite an ∼3-fold difference in P_r_ (Lu et al. 2016; Newman et al. 2017). Accordingly, our correlative functional imaging confirms that the same number of channels can support greater release at these high-P_r_ AZs (Fig. 1). While two-fold greater Ca^2+^ influx at high-P_r_ type Is AZs can explain much of this difference (He et al. 2023; Lu et al. 2016), mounting evidence suggests that here and in other systems differences in P_r_ are separable from Ca^2+^ influx. At distinct inputs formed by CA1 pyramidal neurons, Ca^2+^ influx is greater at high-Pr synapses, but doesn’t explain differences in synaptic strength as raising Ca^2+^ entry at low-Pr synapses to high-Pr synapse levels was not sufficient to increase synaptic strength to high-Pr input levels (Aldahabi et al., 2022). Similar findings have been reported at tonic and phasic synapses of the Crayfish NMJ (Msghina, 1999).

The separability of VGCC abundance, Ca^2+^ influx, and P_r_ appears to be due to molecular and spatial differences between synaptic subtypes. In CA1 pyramidal neurons, differences in Munc-13-dependent SV priming are proposed to establish synapse-specific release properties, possibly due to the presence of distinct isoforms at low- vs. high-P_r_ connections. In the cerebellum, fewer VGCC are more tightly coupled to SVs at high-P_r_ stellate synapses (Rebola et al. 2019). More densely organized VGCCs at the mature vs. developing calyx of Held also exhibit greater coupling with SVs (Chen et al. 2015; Fedchyshyn and Wang 2005; Fekete et al. 2019; Nakamura et al. 2015; Sheng et al. 2012). We find that Cac clusters are denser at high-P_r_ AZs formed by *Drosophila* phasic motor neurons (Fig. 2). Brp, which organizes both VGCCs and SVs, is also more densely organized at type Is synapses (He et al. 2023; Jetti et al. 2023; Mrestani et al. 2021), consistent with an overall more compact organization of high-P_r_ AZs. A straightforward prediction is that a more compact AZ organization will decrease the distance between VGCCs and SVs. Indeed, a recent electrophysiological study using new tools for genetically isolating type Ib and Is inputs demonstrated that neurotransmitter release at denser type Is synapses is less impacted by the slow Ca^2+^ chelator EGTA than type Ib synapses, indicating tighter VGCC-SV coupling (He et al. 2023). Together, these findings suggest that more compact AZs may be a common organizing principle of high-P_r_ synapses. Consistent with this model, Brp and Unc-13 density increase during PHP when P_r_ is increased (Dannhauser et al. 2022; Ghelani et al. 2023; Mrestani et al. 2021).

We also observe molecular differences between *Drosophila* motor inputs that may contribute to distinct P_r_. Somewhat counterintuitively, high-P_r_ type Is AZs have lower levels of both Brp and Stj/α2δ-3 (Figs. 3, 6). Brp/CAST/ELKS AZ cytomatrix proteins are central regulators of synapse organization across species (Dai et al. 2006; Dong et al. 2018; Hallermann et al. 2010; Held, Liu, and Kaeser 2016; Kittel et al. 2006; Liu et al. 2014; McDonald, Fetter, and Shen 2020; Radulovic et al. 2020). Brp interacts broadly with AZ proteins to promote Cac clustering, organize SVs, and recruit Unc13A, which defines SV release sites (Bohme et al. 2016; Fulterer et al. 2018; Ghelani et al. 2023; Liu et al. 2011). While Brp clearly plays a central role in AZ organization and reorganization, we also find that type Ib and Is synapses have distinct requirements for Brp during synapse formation and homeostatic potentiation (Figs. 3, 4). Synapse-specific roles for Brp are supported by a recent functional imaging study in *Drosophila* (Jetti et al. 2023) and studies of ELKS at mammalian inhibitory and excitatory synapses (Held, Liu, and Kaeser 2016), and suggest additional factors establish neuron-specific differences in Brp dependence. A recent single-cell transcriptomic study of type Ib and Is motor neurons provides an unbiased starting point for identifying candidate regulators of molecular differences at low- and high-P_r_ AZs (Jetti et al. 2023). Differentially expressed genes encode cytoskeletal and motor-related proteins, regulators of proteostasis, and post-translational modifying enzymes/pathway components – all of which could potentially contribute to establishing the observed molecular and/or spatial differences between type Ib and Is AZs.

The VGCC complex itself provides an additional potential mechanism for diversifying synaptic function. Both β and α2δ subunits can influence the membrane localization and function of VGCCs (Campiglio and Flucher 2015; Dolphin and Lee 2020; Weiss and Zamponi 2017). In addition to the ability to mix and match subunits, many of the genes encoding VGCC subunits across species are extensively alternatively spliced to generate additional functional diversity (Cingolani et al. 2023; Lipscombe, Andrade, and Allen 2013; Lipscombe and Lopez Soto 2019) – an area of great interest for further investigation at the *Drosophila* NMJ. Because both β and α2δ subunits are generally considered positive regulators of channel trafficking and function, we were surprised to find upon endogenously tagging Stj/α2δ-3 that its levels are lower at high-P_r_ AZs (Figs. 5, 6). Since AZ levels of α subunit Cac are similar at the two inputs, lower levels of Stj at type Is synapses indicates a difference in α:α2δ-3 stoichiometry between low- and high-P_r_ synaptic subtypes. While some previous studies have observed a tight association between the two subunits, a single-molecule tracking and modeling study of mammalian VGCCs found that α2δ subunits have a relatively low affinity for α subunits and predicted a population of α subunits not associated with an α2δ subunit (Cassidy et al. 2014; Voigt et al. 2016).

Consistently, whereas α and β subunits were isolated at near equimolar ratios following affinity purification of Ca_v_2 channels, molar levels of α2δ were a surprising 90% lower (Muller et al. 2010). Our findings suggest there may be a pool of VGCCs lacking an α2δ subunit at endogenous synapses, and further suggest that this pool is specific to or enriched at high-P_r_ type Is AZs. Stj is both required (Dickman, Kurshan, and Schwarz 2008; Kurshan, Oztan, and Schwarz 2009; Ly et al. 2008) and rate limiting (Cunningham et al. 2022) for Cac accumulation at AZs. Stj does not appear to function in the stabilization of channels at the AZ membrane, but rather at an upstream step in the progression from ER to plasma membrane (Cunningham et al. 2022), so complexes may not need to be maintained. Tools for following Stj dynamics in developing neurons will help clarify its precise role in Cac delivery at type Ib and Is AZs.

How might a higher α:α2δ-3 ratio result in higher P_r_? One possibility involves α2δ-3 interactions with cell adhesion molecules. In mammals, α-Neurexin specifically inhibits Ca^2+^ currents in Ca_v_2.2 channels containing α2δ-3 (Tong et al. 2017), which, if similar at the *Drosophila* NMJ, could result in greater inhibition of channel function at type Ib AZs and contribute to the observed difference in Ca^2+^ influx. Loss of *Drosophila* Neurexin reduces neurotransmitter release at the NMJ, but it also leads to reduced synaptogenesis so whether this is a direct effect remains unknown (Li et al. 2007). Another possibility is that Stj isoforms with different functions are differentially expressed at type Ib and Is AZs. A recent transcriptomic study observed no significant differences in mRNA splice isoforms, but transcript levels do not always reflect protein levels (Jetti et al. 2023). Notably, we find that among synapses of either low- or high-P_r_ AZs, Stj levels correlate with P_r_ and Stj levels are increased across AZs of both inputs during acute and chronic PHP (Fig. 7). Cac and Stj levels increase by similar amounts at the two inputs, suggesting the difference in stoichiometry is maintained following homeostatic potentiation. α2δ-3 is an important drug target for treating epilepsy, neuropathic pain, and anxiety, so understanding its cell-specific roles and how α:α2δ-3 stoichiometry impacts channel function and contributes to synaptic diversity is of great interest.

## MATERIALS AND METHODS

### Drosophila genetics and gene editing

The following fly lines used in this study were obtained from the Bloomington Drosophila Stock Center (BDSC, NIH P40OD018537): *w^1118^*(RRID:BDSC_5905), *vasa-Cas9* (RRID:BDSC_51324), piggyBac transposase (RRID:BDSC_8283), and *Df(2R)brp^6.1^* (Gratz et al. 2014; Horn et al. 2003). *brp^69^* and *GluRIIA^sp16^* (*GluRIIA^-/-^*) alleles were generously provided by Stephan Sigrist (Freie Universität Berlin; (Fouquet et al. 2009; Kittel et al. 2006)) and Aaron DiAntonio (Petersen et al. 1997) respectively. *brp* loss-of-function experiments were performed in *brp^69^*/*Df(2R)brp^6.1^*. *Drosophila melanogaster* stocks were raised on molasses food (Lab Express, Type R) in a 25°C incubator with controlled humidity and 12h light/dark cycle. Endogenously tagged *cac*, *Ca-β*, *stolid*, and *straightjacket* (*stj*) alleles were generated using our piggyBac-based CRISPR approach as previously detailed (flyCRISPR.molbio.wisc.edu; (Bruckner et al. 2017; Gratz et al. 2019)). We used FlyBase (release FB2024_01) to obtain genomic sequences (doi.org/10.1093/genetics/iyad211). Td-Tomato and Halo tags were incorporated at the N-terminus of Cac, where we have previously incorporated fluorescent tags (Ghelani et al. 2023; Gratz et al. 2019). V5 tags were incorporated at the N-terminus of Stj and Stolid after their signal peptide sequences, immediately after Asparagine 34 of Stj and after Isoleucine 34 of Stolid. For Ca-β, V5 was inserted immediately after Proline 446 of isoform Ca-β-PN. All endogenously tagged lines are fully viable in homozygous males and females and were molecularly confirmed by Sanger sequencing.

### Immunostaining

All antibodies used, associated fixation methods, and incubation times can be found in Table S2. Male wandering third-instar larvae were dissected in ice-cold saline and fixed either for 6 minutes at room temperature with Bouin’s fixative, 5 minutes on ice with 100% methanol, or 30 minutes at room temperature (RT) in 4% PFA. Dissections were permeabilized with PTX (PBS with 0.1% Triton-X 100) and blocked for 1 hour at RT using 5% goat serum and 1% bovine serum albumin. Stained larvae were mounted in Vectashield (Vector Laboratories, #H-1000) under Fisherbrand coverglass (Fisher Scientific, #12541B) for confocal microscopy, with Prolong glass mounting medium (ThermoFisher Scientific, #P36980) under Zeiss High Performance Coverglass (Zeiss, #474030-9000-000) for super-resolution optical reassignment microscopy, or buffer (see STORM imaging and analysis section) under Zeiss coverglass with edges sealed using vacuum grease for STORM microscopy.

### Ca^2+^ imaging and analysis

Functional imaging was performed on a Nikon A1R resonant scanning confocal mounted on a FN1 microscope using a Nikon Apo LWD 25x 1.1 NA objective and a Mad City Labs piezo drive nosepiece. Dissections and data collection were performed as previously described in Gratz et al., 2019. Briefly, c*ac^Td-Tomato-N^*; *Mhc-GCaMP6f* male 3rd instar larvae were dissected in HL3 containing 0.2mM Ca^2+^ and 25mM Mg^2+^ with motor axons severed and the larval brain removed. Larval filets were placed in HL3 containing 1.5mM Ca^2+^ and 25mM Mg^2+^ for recording. Nerves were suctioned into 1.5mm pipettes and stimulus amplitude was adjusted to recruit both type Ib and Is inputs. Motor terminals from segments A2-4 at NMJ 6/7 were imaged for Cac^Td-Tomato-N^ levels first using a galvanometer scanner, then a resonant scanner to collect GCaMP6f events in a single focal plane continuously for 120 stimulations at a 0.2 Hz stimulation frequency.

Z-stacks and movies were loaded into Nikon Elements Software (NIS) where movies were motion corrected, background subtracted, and denoised. Change in fluorescence (ΔF) movies were then created by subtracting the average of the previous 10 frames from each frame. A substack of only stimulation frames was further processed using a gaussian filter followed by the Bright Spots detection module in the Nikon GA3 software to identify the location of each postsynaptic event. Cac^Td-Tomato-N^ fluorescence intensity levels and coordinate locations were measured for 531 AZs for type Ib and 365 AZs for type Is terminals across 6 animals. X-Y coordinate positions of fluorescent signals from GCaMP6f postsynaptic events were aligned to Cac^Td-Tomato-N^ puncta locations and each post synaptic event assigned to a Cac punctum using nearest neighbor analysis. Postsynaptic events that did not map within 960 nm of a Cac^Td-Tomato-N^ punctum were discarded from the analysis. Pearson’s correlation was used to determine the correlation between P_r_ and Cac levels normalized to average to account for variability between imaging sessions. Cac intensity-P_r_ heat maps were generated using Python matplotlib and seaborn plotting packages.

### STORM imaging and analysis

STORM imaging was performed on a Nikon Eclipse Ti2 3D NSTORM with an Andor iXon Ultra camera, Nikon LUN-F 405/488/640 nm lasers, and a Nikon 100x 1.49 NA objective. STORM buffer (10mM MEA (pH 8.0), 3 U/mL pyranose oxidase, and 90 U/mL catalase, 10% (w/v) glucose, 10 mM sodium chloride, and 50 mM Tris hydrochloride) was made fresh each imaging day and pH adjusted to between 7.0-8.0 using acetic acid. *cac^HaloTag-N^* NMJs were labeled as detailed in Table S2 and immediately imaged for HRP in 488 channel to identify type Ib and Is terminals. Cac^HaloTag-N^ was then imaged using the 640 nm laser line at 33 Hz for 5000 frames. 405 nm laser power was gradually increased over the course of imaging to compensate for run-down of blinking rates. A back aperture camera was used to ensure beam focus and position for each imaging session to ensure high signal to noise. Data were binned with a CCD minimum threshold of 100 and drift correction was applied using the NIS Software STORM package. ROIs of single boutons were drawn in NIS using HRP in the 488 nm channel followed by a DBSCAN analysis with criteria of 10 molecules within 50 nm to determine clusters. Positional coordinates of localizations within clusters from DBSCAN were exported from NIS and run through a Python script published with this manuscript. Using the implementation developed in Mrestani et al., 2021 as a starting point, we wrote custom code to use the alpha shapes component of the CGAL package (https://www.cgal.org), via a python wrapper (https://anaconda.org/conda-forge/cgal), to measure the area of Ca^2+^ channel clusters, the number of localizations, and calculate cluster density. To achieve an average lateral localization accuracy of ∼30 nm, all localizations with >50 nm localization accuracy were removed prior to analysis. Using this custom code, Cac^HaloTag-N^ area was analyzed using an alpha value of 0.015, which controls the complexity of cluster boundaries (not restricted to be convex).

### Confocal imaging and analysis

For quantitative AZ analysis of larval NMJs, dissections stained in the same dish were imaged on a Nikon Eclipse Ni A1R+ confocal microscope using an Apo TIRF 60x 1.49 NA oil-immersion objective for larval NMJs. NMJs containing both type Is and type Ib branches from muscles 6/7 in segments A2-4 were collected. ROIs were drawn using HRP staining to differentiate between type Ib and Is branches. To analyze individual AZs, Nikon Elements GA3 Software was used to process images with Gaussian and rolling ball filters and measure fluorescence intensity levels at individual puncta identified by the Bright Spots module. When experimental design allowed, Brp fluorescence signal was used to create a binary mask to aid in the identification of AZ ROIs for analysis. Otherwise, binary masks were created based on the fluorescence signal of the channel analyzed. Confocal fluorescence intensity level data are reported as the sum fluorescence intensity per AZ averaged over individual NMJs. For Fig. 5, larvae were stained separately and imaged using a Nikon Plan-Apo 20x 0.75 NA objective (ventral ganglia) or Apo TIRF 60x 1.49 NA oil-immersion objective (NMJs).

Super-resolution optical reassignment images were obtained on a Nikon CSU-W1 SoRa (Spinning Disk Super Resolution by Optical Pixel Reassignment) with a Photometrics Prime BSI sCMOS camera and a 60x 1.49 NA oil-immersion objective. Images were acquired using Nikon NIS and deconvolved using Richardson-Lucy deconvolution with 15-20 iterations.

### Electrophysiology

Current-clamp recordings were performed as previously described (Bruckner et al. 2017). Male third-instar larvae were dissected in HL3 (70 mM NaCl, 5 mM KCl, 15 mM MgCl2, 10 mM NaHCO3, 115 mM sucrose, 5 mM trehalose, 5 mM HEPES, pH 7.2) with 0.25 mM Ca^2+^. Recordings were performed in HL3 at the external Ca^2+^ concentration indicated. Sharp borosilicate electrodes filled with 3 M KCl were used to record from muscle 6 of abdominal segments A3 and A4. Recordings were conducted on a Nikon FN1 microscope using a 40x 0.80 NA water-dipping objective and acquired using an Axoclamp 900A amplifier, Digidata 1550B acquisition system, and pClamp 11.0.3 software (Molecular Devices).

For each cell with an initial resting potential between −60 and −80 mV and input resistance ≥5 MΩ, mean miniature excitatory junctional potentials (mEJPs) were collected for 1 minute in the absence of stimulation and analyzed using Mini Analysis (Synaptosoft). EJPs were generated by applying a stimulus to severed segmental nerves at a frequency of 0.2 Hz using an isolated pulse stimulator 2100 (A-M Systems). Stimulus amplitude was adjusted to consistently elicit compound responses from both type Ib and Is motor neurons. At least 25 consecutive EJPs were recorded for each cell and analyzed in pClamp to obtain mean amplitude. Quantal content was calculated for each recording as mean EJP amplitude divided by mean mEJP amplitude.

### Acute homeostatic challenge

Acute PHP was induced by incubating semi-intact preparations in 20 µM Philanthotoxin-433 (Fig. 4: PhTx; Santa Cruz, sc-255421, Lot B1417 and Fig. 7: PhTx; Sigma Aldrich, P207-2, Lot MKCK7405) diluted in HL3 containing 0.4 mM Ca^2+^ for 10 minutes at room temperature (Frank et al. 2006). Control preparations were given a mock treatment. Following control and experimental treatment, dissections were completed, fixed in 4% PFA for 30 minutes (*cac^sfGFP-N^*) or 100% ice-cold methanol on ice for 5 minutes (*stj^V5-N^*), and stained in the same dish. Analyses of fluorescent intensity levels were performed as previously described in the *Confocal imaging and analysis section*.

### Experimental design and statistical analysis

Statistical analyses were conducted in GraphPad Prism 9. Normality was determined by the D’Agostino–Pearson omnibus test. Comparisons of normally distributed data were conducted by Student’s *t* test (with Welch’s correction in the case of unequal variance) for single comparisons and ANOVA followed by Tukey’s test for multiple comparisons. For non-normally distributed data, the Mann–Whitney *U* test and Kruskal-Wallis test followed by Dunn’s multiple comparisons tests were used for single and multiple comparisons, respectively. Paired analysis of non-normally distributed data was conducted using Wilcoxon’s matched-pairs signed rank test. One-dimensional Pearson correlation coefficients (*r*) were used to compare intensity levels and neurotransmitter release probability. ANCOVA test was performed on all regression lines to determine if slopes were significantly different. Reported values are mean ± SEM. Sample size, statistical test, and *p* values for each comparison are reported in Table S1.

## ACKNOWLEDGEMENTS

We thank the Developmental Studies Hybridoma Bank, the Bloomington *Drosophila* Stock Center (NIH P40OD018537), Flybase (Öztürk-Çolak et al. 2024), Ehud Isacoff (UC Berkeley), and Stephan Sigrist (Freie Universität Berlin) for providing antibodies and fly stocks. The Nikon SoRa/STORM microscope was generously provided by The Neurobiology of Cells and Circuits/Center for Translational Neuroscience Microscopy Committee, Carney Institute for Brain Science. We are grateful to Joel Hirsch (Tel Aviv University) for consultations on tagging Ca-β, Nicholas Deakin (Nikon) for guidance on STORM imaging, the Heckmann lab (University of Würzburg) for guidance on STORM image analysis pipelines, Matthew Knoeppel for help generating CRISPR alleles, and Liana Lewis for her assistance with image analysis. We thank Rajan Thakur and the members of the O’Connor-Giles lab for thoughtful discussions and comments on the manuscript. This work was supported by grants from the National Institute of Neurological Disorders and Stroke, National Institutes of Health to K.M.O.G (R01NS078179) and Audrey Medeiros (F31NS122424), Brown Neuroscience Graduate Program training grant T32 MH020068, and funds from the Brown University Carney Institute for Brain Science.

---

**Table S1.**
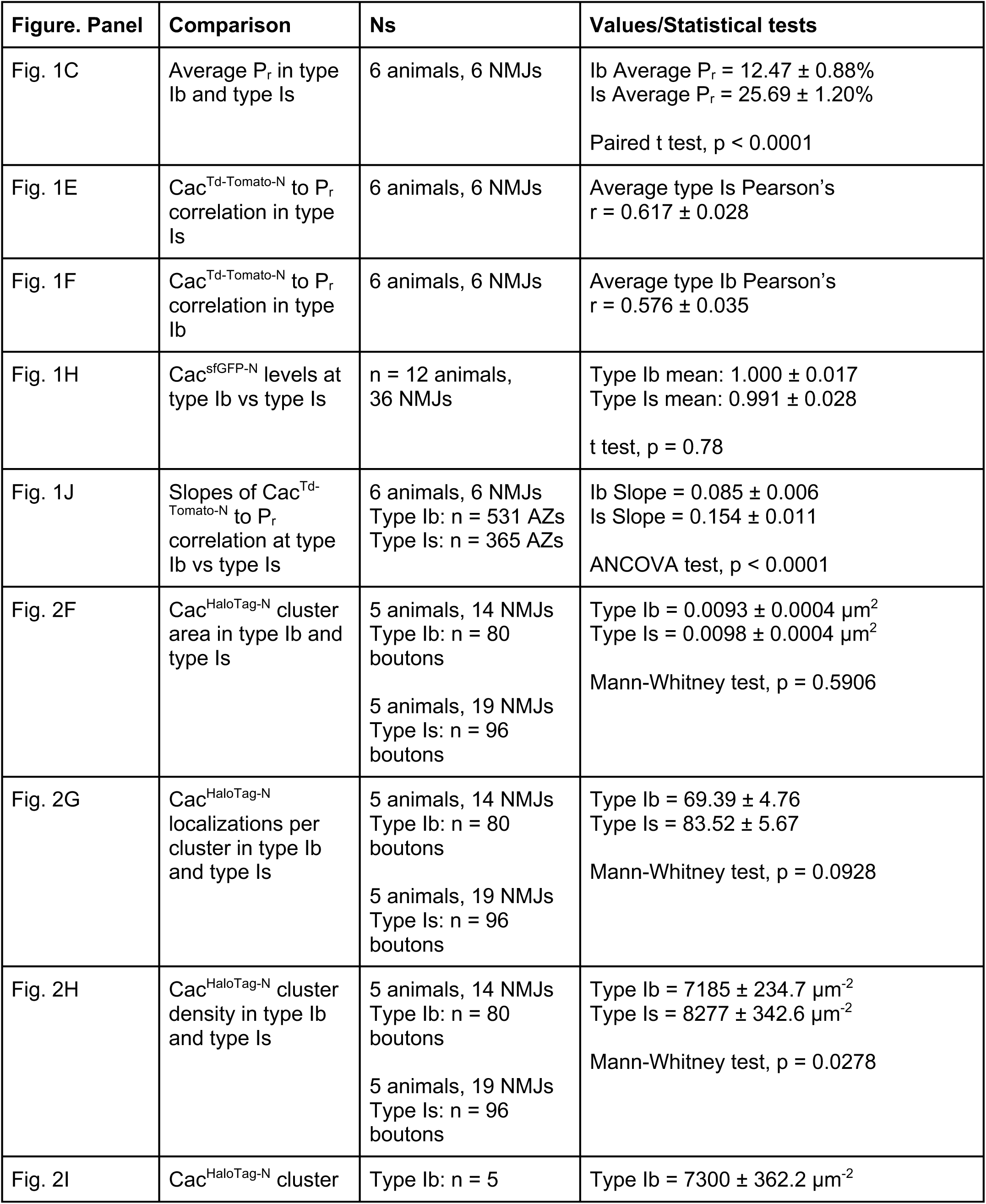
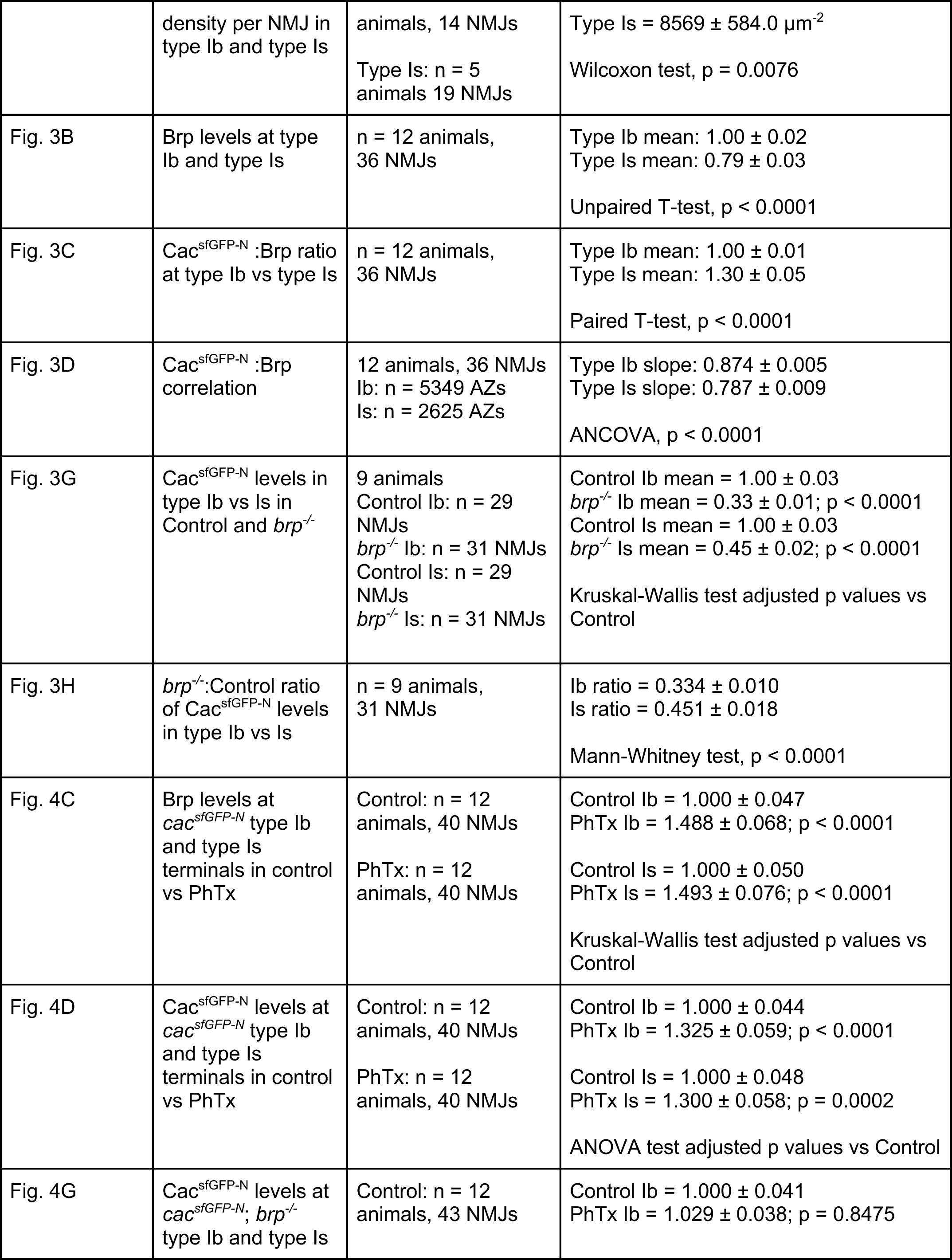
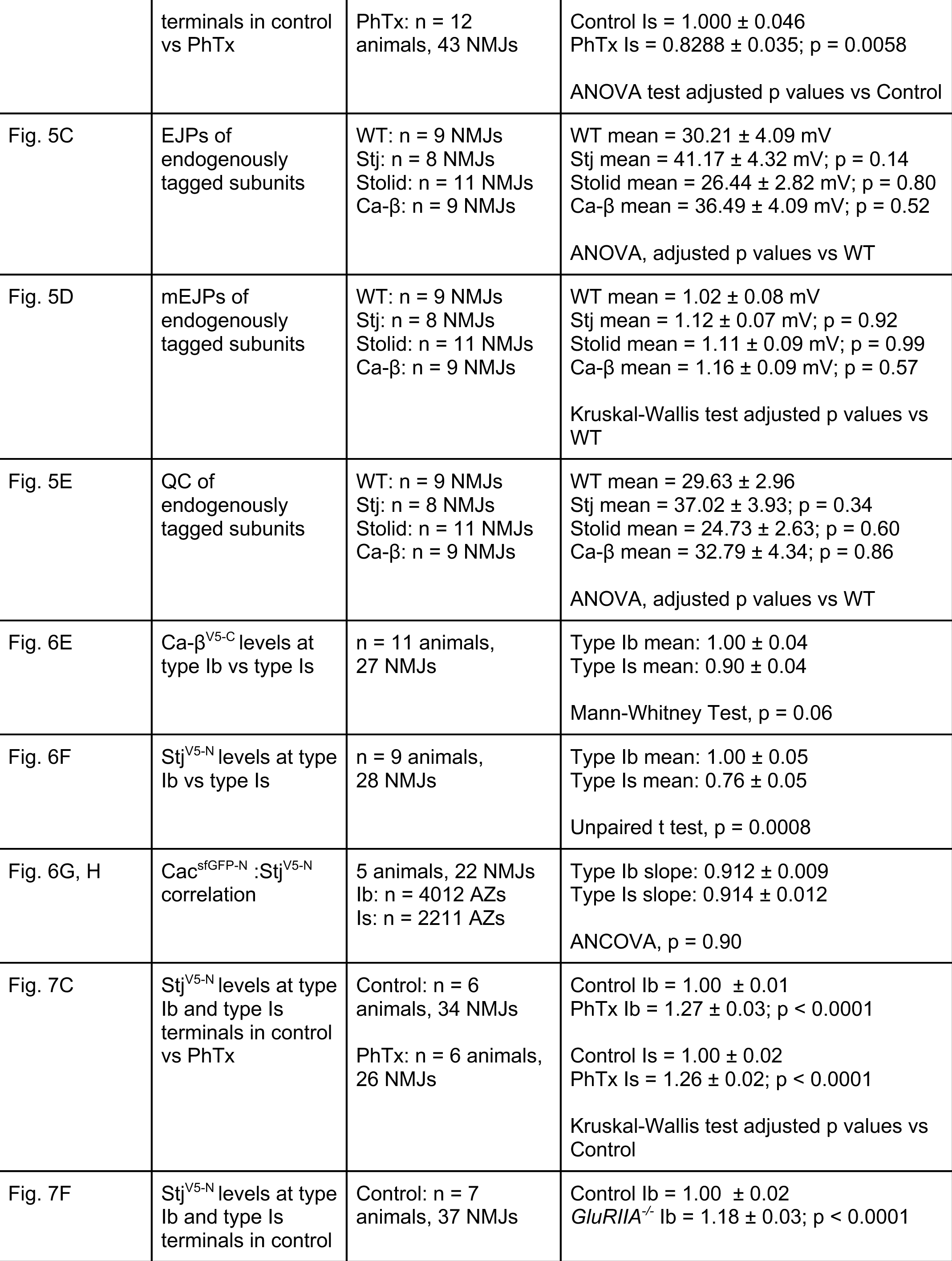
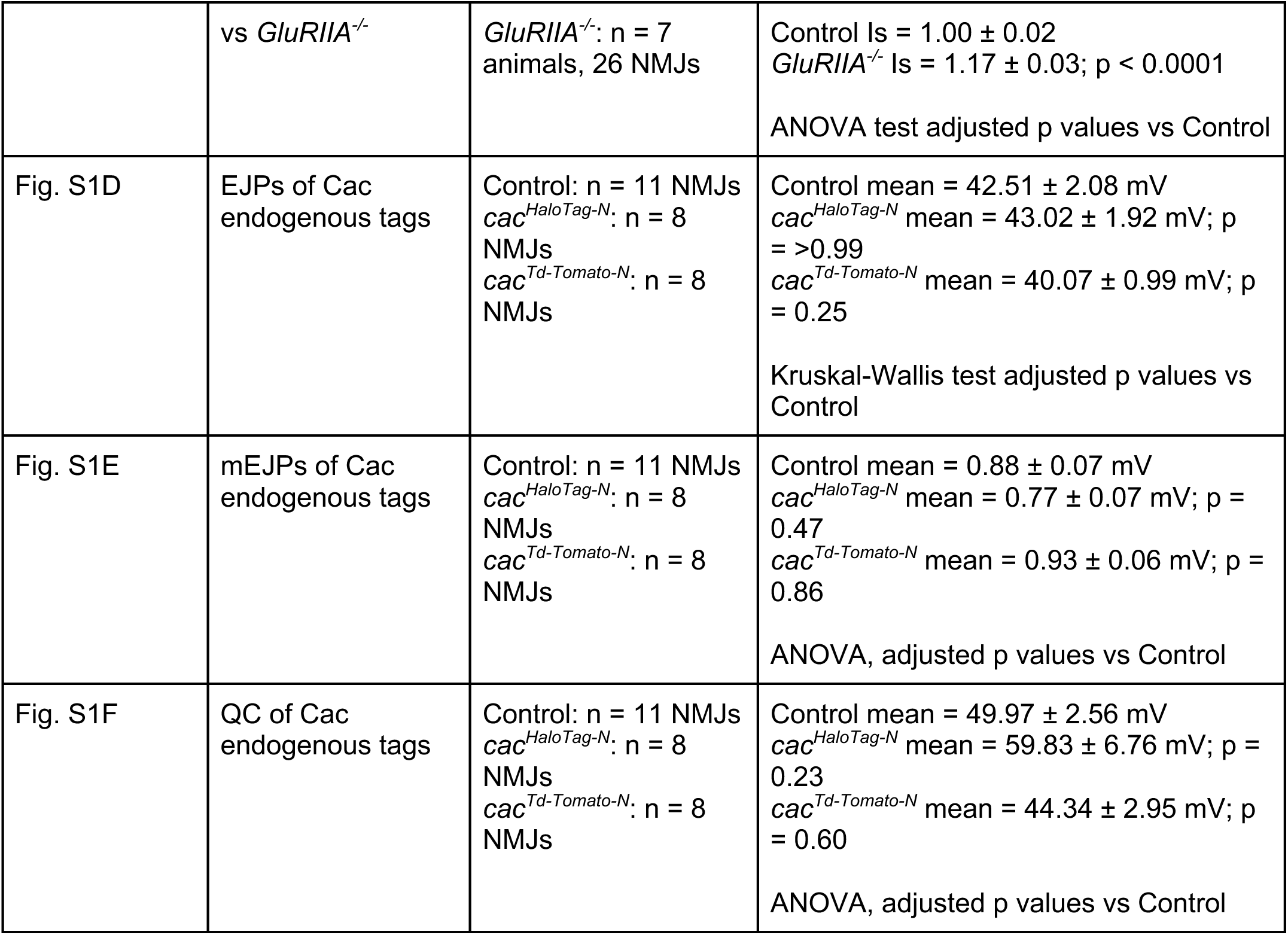
Absolute values and statistics. All comparisons, Ns (animals, NMJs, AZs (AZs), and statistical tests used in this study. All values are mean ± SEM.

**Table S2.**
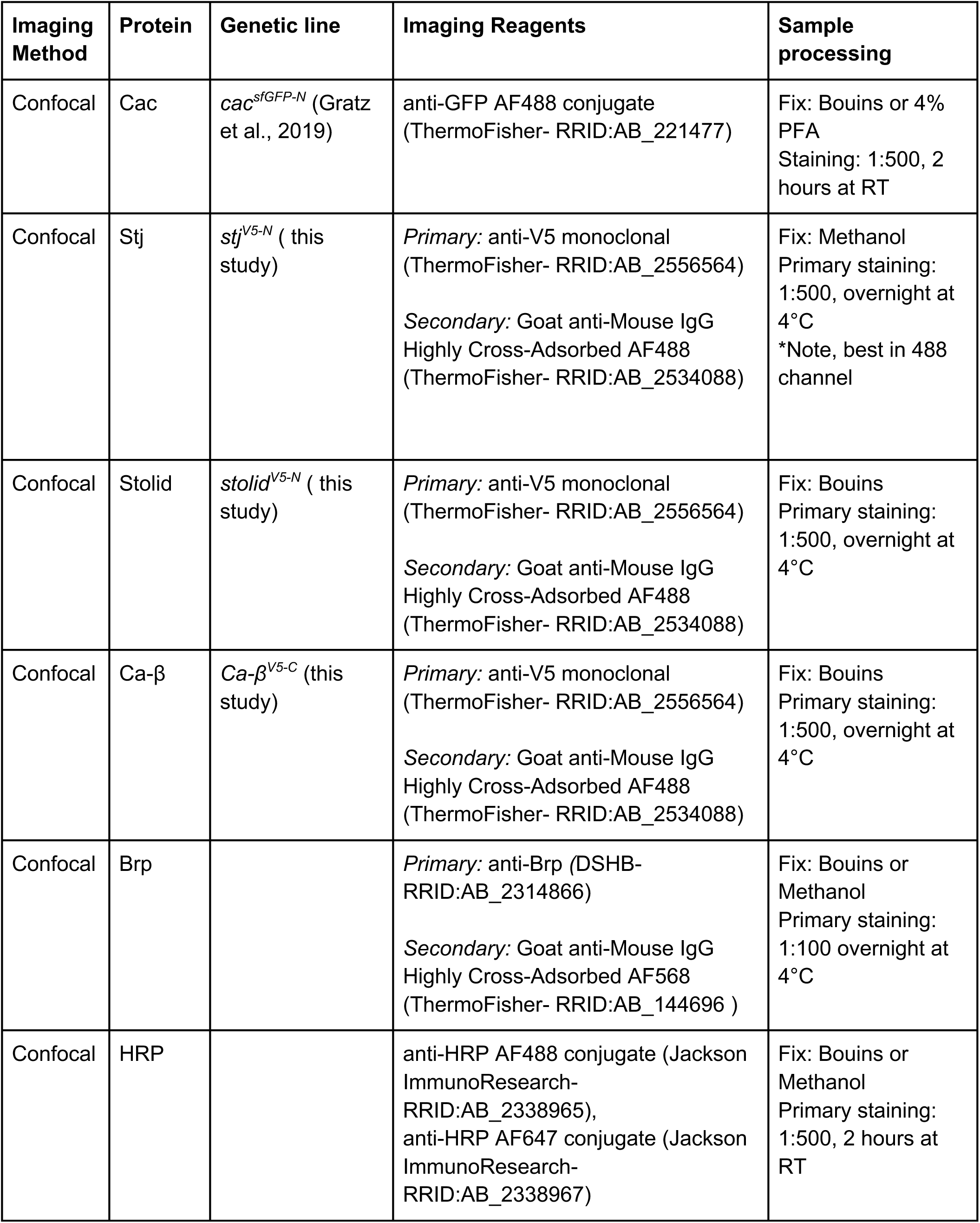
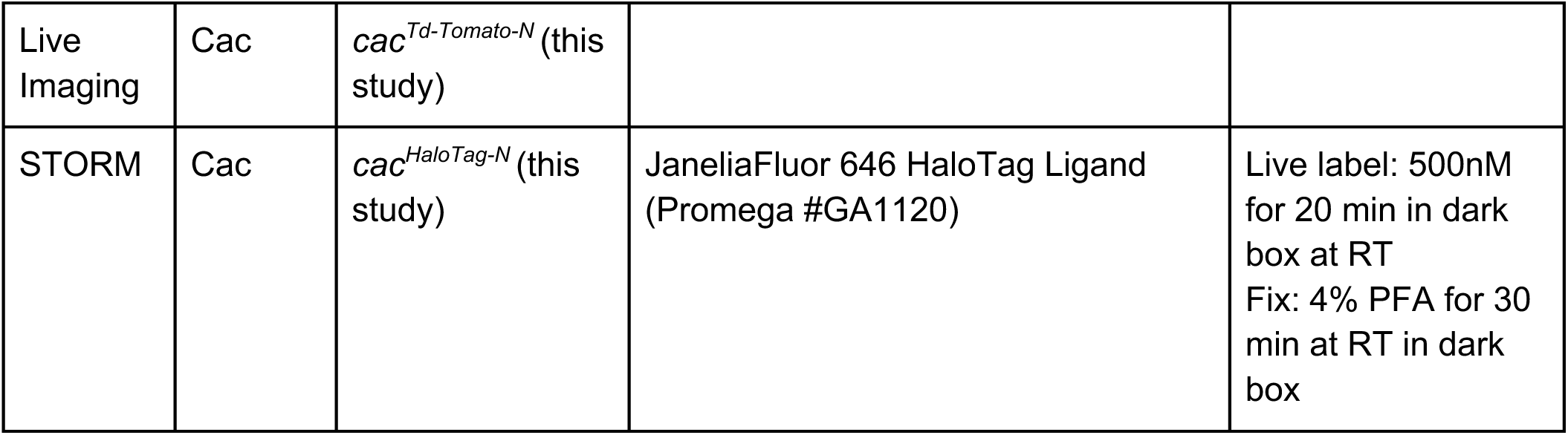
Imaging details. This table contains detailed information on how each protein was labeled and visualized using live and fixed confocal microscopy and STORM imaging. All secondary antibodies were incubated at RT for 2 hours at a concentration of 1:500.

**Figure S1:**
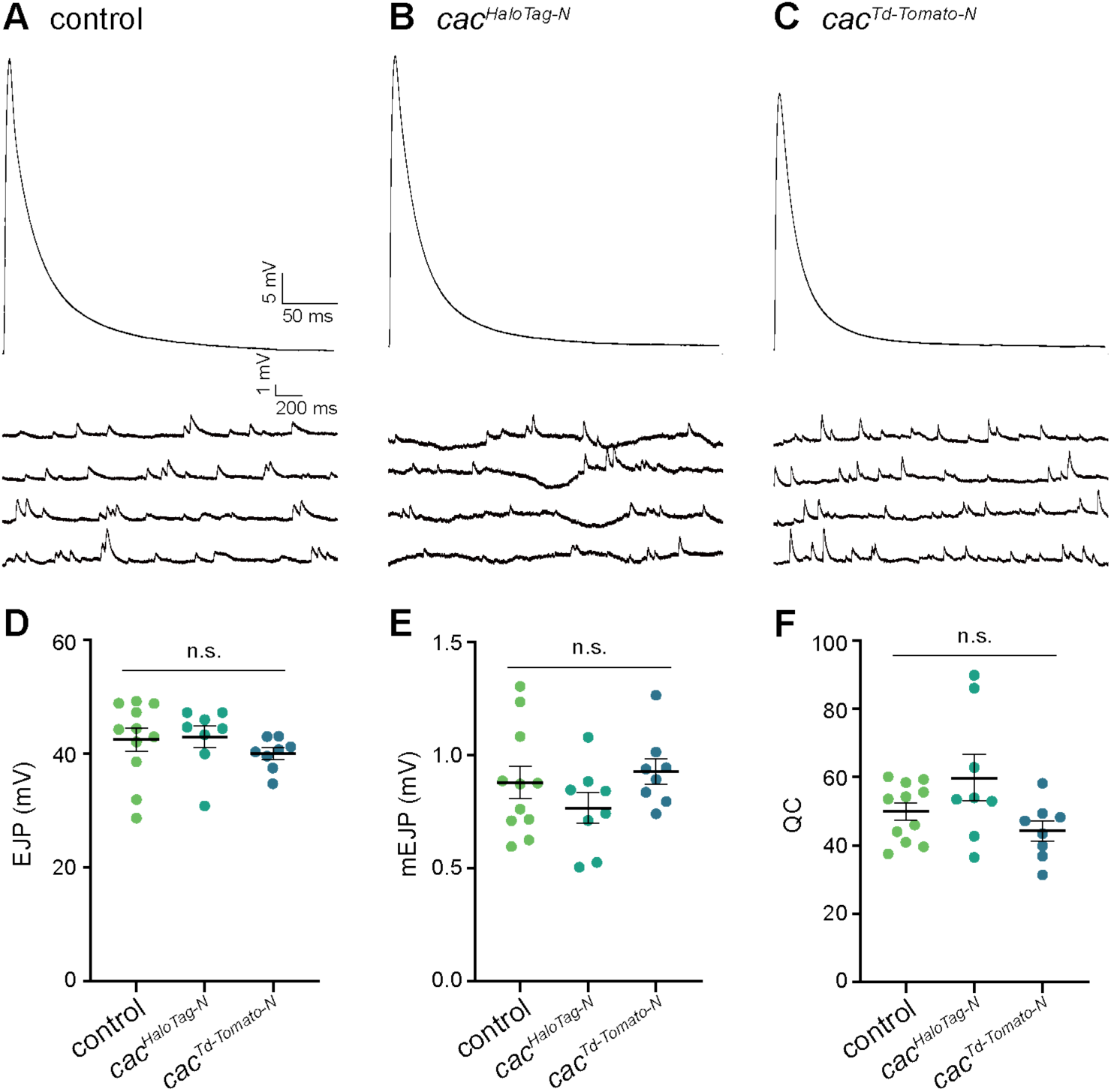
Electrophysiological validation of endogenously tagged cacophony lines. **(A-C)** Representative traces of EJPs (top) and mEJPs (bottom) in control, c*ac^HaloTag-N^*, and *cac^Td-Tomato-N^*. **(D-F)** Quantification of EJPs, mEJPs, and quantal content (QC).

## REFERENCES

Akbergenova, Yulia, Karen L. Cunningham, Yao V. Zhang, Shirley Weiss, and J. Troy Littleton. 2018. ’Characterization of developmental and molecular factors underlying release heterogeneity at Drosophila synapses’, eLife, 7.

Aldahabi, M., F. Balint, N. Holderith, A. Lorincz, M. Reva, and Z. Nusser. 2022. ’Different priming states of synaptic vesicles underlie distinct release probabilities at hippocampal excitatory synapses’, Neuron, 110: 4144–61 e7.

Aponte-Santiago, N. A., and J. T. Littleton. 2020. ’Synaptic Properties and Plasticity Mechanisms of Invertebrate Tonic and Phasic Neurons’, Front Physiol, 11: 611982.

Aponte-Santiago, N. A., K. G. Ormerod, Y. Akbergenova, and J. T. Littleton. 2020. ’Synaptic Plasticity Induced by Differential Manipulation of Tonic and Phasic Motoneurons in Drosophila’, J Neurosci, 40: 6270–88.

Ariel, P., M. B. Hoppa, and T. A. Ryan. 2012. ’Intrinsic variability in Pv, RRP size, Ca(2+) channel repertoire, and presynaptic potentiation in individual synaptic boutons’, Front Synaptic Neurosci, 4: 9.

Atwood, H. L., C. K. Govind, and C.-F. Wu. 1993. ’Differential ultrastructure of synaptic terminals on ventral longitudinal abdominal muscles in Drosophila larvae’, Journal of Neurobiology, 24: 1008–24.

Atwood, H. L., and S. Karunanithi. 2002. ’Diversification of synaptic strength: presynaptic elements’, Nat Rev Neurosci, 3: 497–516.

Bauer, C. S., A. Tran-Van-Minh, I. Kadurin, and A. C. Dolphin. 2010. ’A new look at calcium channel alpha2delta subunits’, Curr Opin Neurobiol, 20: 563–71.

Bohme, M. A., C. Beis, S. Reddy-Alla, E. Reynolds, M. M. Mampell, A. T. Grasskamp, J. Lutzkendorf, D. D. Bergeron, J. H. Driller, H. Babikir, F. Gottfert, I. M. Robinson, C. J. O’Kane, S. W. Hell, M. C. Wahl, U. Stelzl, B. Loll, A. M. Walter, and S. J. Sigrist. 2016. ’Active zone scaffolds differentially accumulate Unc13 isoforms to tune Ca(2+) channel-vesicle coupling’, Nat Neurosci, 19: 1311–20.

Bohme, M. A., A. W. McCarthy, A. T. Grasskamp, C. B. Beuschel, P. Goel, M. Jusyte, D. Laber, S. Huang, U. Rey, A. G. Petzoldt, M. Lehmann, F. Gottfert, P. Haghighi, S. W. Hell, D. Owald, D. Dickman, S. J. Sigrist, and A. M. Walter. 2019. ’Rapid active zone remodeling consolidates presynaptic potentiation’, Nat Commun, 10: 1085.

Branco, T., and K. Staras. 2009. ’The probability of neurotransmitter release: variability and feedback control at single synapses’, Nat Rev Neurosci, 10: 373–83.

Bruckner, J. J., H. Zhan, S. J. Gratz, M. Rao, F. Ukken, G. Zilberg, and K. M. O’Connor-Giles. 2017. ’Fife organizes synaptic vesicles and calcium channels for high-probability neurotransmitter release’, J Cell Biol, 216: 231–46.

Campiglio, Marta, and Bernhard E. Flucher. 2015. ’The Role of Auxiliary Subunits for the Functional Diversity of Voltage-Gated Calcium Channels’, Journal of Cellular Physiology, 230: 2019–31.

Cao, Yu-Qing, Erika S. Piedras-Rentería, Geoffrey B. Smith, Gong Chen, Nobutoshi C. Harata, and Richard W. Tsien. 2004. ’Presynaptic Ca^2+^ Channels Compete for Channel Type-Preferring Slots in Altered Neurotransmission Arising from Ca^2+^ Channelopathy’, Neuron, 43: 387–400.

Cassidy, J. S., L. Ferron, I. Kadurin, W. S. Pratt, and A. C. Dolphin. 2014. ’Functional exofacially tagged N-type calcium channels elucidate the interaction with auxiliary alpha2delta-1 subunits’, Proc Natl Acad Sci U S A, 111: 8979–84.

Chen, Z., B. Das, Y. Nakamura, D. A. DiGregorio, and S. M. Young, Jr. 2015. ’Ca2+ channel to synaptic vesicle distance accounts for the readily releasable pool kinetics at a functionally mature auditory synapse’, J Neurosci, 35: 2083–100.

Cingolani, Lorenzo A., Agnes Thalhammer, Fanny Jaudon, Jessica Muià, and Gabriele Baj. 2023. ’Nanoscale organization of CaV2.1 splice isoforms at presynaptic terminals: implications for synaptic vesicle release and synaptic facilitation’, 404: 931–37.

Cunningham, K. L., C. W. Sauvola, S. Tavana, and J. T. Littleton. 2022. ’Regulation of presynaptic Ca(2+) channel abundance at active zones through a balance of delivery and turnover’, eLife, 11.

Dai, Y., H. Taru, S. L. Deken, B. Grill, B. Ackley, M. L. Nonet, and Y. Jin. 2006. ’SYD-2 Liprin-alpha organizes presynaptic active zone formation through ELKS’, Nat Neurosci, 9: 1479–87.

Dannhauser, S., A. Mrestani, F. Gundelach, M. Pauli, F. Komma, P. Kollmannsberger, M. Sauer, M. Heckmann, and M. M. Paul. 2022. ’Endogenous tagging of Unc-13 reveals nanoscale reorganization at active zones during presynaptic homeostatic potentiation’, Front Cell Neurosci, 16: 1074304.

Davis, G. W., and M. Muller. 2015. ’Homeostatic control of presynaptic neurotransmitter release’, Annu Rev Physiol, 77: 251–70.

Dickman, D. K., P. T. Kurshan, and T. L. Schwarz. 2008. ’Mutations in a Drosophila alpha2delta voltage-gated calcium channel subunit reveal a crucial synaptic function’, J Neurosci, 28: 31–8.

Dolphin, A. C. 2018. ’Voltage-gated calcium channel alpha (2)delta subunits: an assessment of proposed novel roles’, F1000Res, 7.

Dolphin, A. C., and A. Lee. 2020. ’Presynaptic calcium channels: specialized control of synaptic neurotransmitter release’, Nat Rev Neurosci, 21: 213–29.

Dong, W., T. Radulovic, R. O. Goral, C. Thomas, M. Suarez Montesinos, D. Guerrero-Given, A. Hagiwara, T. Putzke, Y. Hida, M. Abe, K. Sakimura, N. Kamasawa, T. Ohtsuka, and S. M. Young, Jr. 2018. ’CAST/ELKS Proteins Control Voltage-Gated Ca(2+) Channel Density and Synaptic Release Probability at a Mammalian Central Synapse’, Cell Rep, 24: 284–93 e6.

Eggermann, E., I. Bucurenciu, S. P. Goswami, and P. Jonas. 2011. ’Nanodomain coupling between Ca(2)(+) channels and sensors of exocytosis at fast mammalian synapses’, Nat Rev Neurosci, 13: 7–21.

Ehmann, Nadine, Sebastian Van De Linde, Amit Alon, Dmitrij Ljaschenko, Xi Zhen Keung, Thorge Holm, Annika Rings, Aaron Diantonio, Stefan Hallermann, Uri Ashery, Manfred Heckmann, Markus Sauer, and Robert J. Kittel. 2014. ’Quantitative super-resolution imaging of Bruchpilot distinguishes active zone states’, Nature Communications, 5.

Fedchyshyn, M. J., and L. Y. Wang. 2005. ’Developmental transformation of the release modality at the calyx of Held synapse’, J Neurosci, 25: 4131–40.

Fekete, A., Y. Nakamura, Y. M. Yang, S. Herlitze, M. D. Mark, D. A. DiGregorio, and L. Y. Wang. 2019. ’Underpinning heterogeneity in synaptic transmission by presynaptic ensembles of distinct morphological modules’, Nat Commun, 10: 826.

Fouquet, W., D. Owald, C. Wichmann, S. Mertel, H. Depner, M. Dyba, S. Hallermann, R. J. Kittel, S. Eimer, and S. J. Sigrist. 2009. ’Maturation of active zone assembly by Drosophila Bruchpilot’, J Cell Biol, 186: 129–45.

Frank, C. A. 2014. ’Homeostatic plasticity at the Drosophila neuromuscular junction’, Neuropharmacology, 78: 63–74.

Frank, C. A., M. J. Kennedy, C. P. Goold, K. W. Marek, and G. W. Davis. 2006. ’Mechanisms underlying the rapid induction and sustained expression of synaptic homeostasis’, Neuron, 52: 663–77.

Früh, Susanna M., Ulf Matti, Philipp R. Spycher, Marina Rubini, Sebastian Lickert, Thomas Schlichthaerle, Ralf Jungmann, Viola Vogel, Jonas Ries, and Ingmar Schoen. 2021. ’Site-Specifically-Labeled Antibodies for Super-Resolution Microscopy Reveal In Situ Linkage Errors’, ACS Nano, 15: 12161–70.

Fulterer, A., T. F. M. Andlauer, A. Ender, M. Maglione, K. Eyring, J. Woitkuhn, M. Lehmann, T. Matkovic-Rachid, J. R. P. Geiger, A. M. Walter, K. I. Nagel, and S. J. Sigrist. 2018. ’Active Zone Scaffold Protein Ratios Tune Functional Diversity across Brain Synapses’, Cell Rep, 23: 1259–74.

Genc, O., and G. W. Davis. 2019. ’Target-wide Induction and Synapse Type-Specific Robustness of Presynaptic Homeostasis’, Curr Biol, 29: 3863–73 e2.

Ghelani, Tina, Marc Escher, Ulrich Thomas, Klara Esch, Janine Lützkendorf, Harald Depner, Marta Maglione, Pierre Parutto, Scott Gratz, Tanja Matkovic-Rachid, Stefanie Ryglewski, Alexander M. Walter, David Holcman, Kate O‘Connor Giles, Martin Heine, and Stephan J. Sigrist. 2023. ’Interactive nanocluster compaction of the ELKS scaffold and Cacophony Ca^2+^ channels drives sustained active zone potentiation’, Science Advances, 9: eade7804.

Gratz, S. J., F. P. Ukken, C. D. Rubinstein, G. Thiede, L. K. Donohue, A. M. Cummings, and K. M. O’Connor-Giles. 2014. ’Highly specific and efficient CRISPR/Cas9-catalyzed homology-directed repair in Drosophila’, Genetics, 196: 961–71.

Gratz, Scott J., Pragya Goel, Joseph J. Bruckner, Roberto X. Hernandez, Karam Khateeb, Gregory T. Macleod, Dion Dickman, and Kate M. O’Connor-Giles. 2019. ’Endogenous tagging reveals differential regulation of Ca^2+^ channels at single AZs during presynaptic homeostatic potentiation and depression’, The Journal of Neuroscience: 3068–18.

Grimm, J. B., B. P. English, J. Chen, J. P. Slaughter, Z. Zhang, A. Revyakin, R. Patel, J. J. Macklin, D. Normanno, R. H. Singer, T. Lionnet, and L. D. Lavis. 2015. ’A general method to improve fluorophores for live-cell and single-molecule microscopy’, Nat Methods, 12: 244–50, 3 p following 50.

Guerrero, G., D. F. Reiff, G. Agarwal, R. W. Ball, A. Borst, C. S. Goodman, and E. Y. Isacoff. 2005. ’Heterogeneity in synaptic transmission along a Drosophila larval motor axon’, Nat Neurosci, 8: 1188–96.

Hallermann, S., R. J. Kittel, C. Wichmann, A. Weyhersmuller, W. Fouquet, S. Mertel, D. Owald, S. Eimer, H. Depner, M. Schwarzel, S. J. Sigrist, and M. Heckmann. 2010. ’Naked dense bodies provoke depression’, J Neurosci, 30: 14340–5.

Hatt, H., and D. O. Smith. 1976. ’Non-uniform probabilities of quantal release at the crayfish neuromuscular junction’, J Physiol, 259: 395–404.

He, K., Y. Han, X. Li, R. X. Hernandez, D. V. Riboul, T. Feghhi, K. A. Justs, O. Mahneva, S. Perry, G. T. Macleod, and D. Dickman. 2023. ’Physiologic and Nanoscale Distinctions Define Glutamatergic Synapses in Tonic vs Phasic Neurons’, J Neurosci, 43: 4598–611.

Heinrich, Laurin, and Stefanie Ryglewski. 2020. ’Different functions of two putative Drosophila α2δ subunits in the same identified motoneurons’, Scientific Reports, 10: 13670.

Held, R. G., C. Liu, and P. S. Kaeser. 2016. ’ELKS controls the pool of readily releasable vesicles at excitatory synapses through its N-terminal coiled-coil domains’, eLife, 5.

Holderith, Noemi, Andrea Lorincz, Gergely Katona, Balázs Rózsa, Akos Kulik, Masahiko Watanabe, and Zoltan Nusser. 2012. ’Release probability of hippocampal glutamatergic terminals scales with the size of the active zone’, Nat Neurosci, 15: 988–97.

Hoover, K. M., S. J. Gratz, N. Qi, K. A. Herrmann, Y. Liu, J. J. Perry-Richardson, P. J. Vanderzalm, K. M. O’Connor-Giles, and H. T. Broihier. 2019. ’The calcium channel subunit alpha(2)delta-3 organizes synapses via an activity-dependent and autocrine BMP signaling pathway’, Nat Commun, 10: 5575.

Hoppa, Michael B., Beatrice Lana, Wojciech Margas, Annette C. Dolphin, and Timothy A. Ryan. 2012. ’α2δ expression sets presynaptic calcium channel abundance and release probability’, Nature, 486: 122–25.

Horn, C., N. Offen, S. Nystedt, U. Hacker, and E. A. Wimmer. 2003. ’piggyBac-based insertional mutagenesis and enhancer detection as a tool for functional insect genomics’, Genetics, 163: 647–61.

James, T. D., D. J. Zwiefelhofer, and C. A. Frank. 2019. ’Maintenance of homeostatic plasticity at the Drosophila neuromuscular synapse requires continuous IP(3)-directed signaling’, eLife, 8.

Jetti, Suresh K., Andrés B. Crane, Yulia Akbergenova, Nicole A. Aponte-Santiago, Karen L. Cunningham, Charles A. Whittaker, and J. Troy Littleton. 2023. ’Molecular logic of synaptic diversity between Drosophila tonic and phasic motoneurons’, Neuron, 111: 3554–69.e7.

Kanamori, T., M. I. Kanai, Y. Dairyo, K. Yasunaga, R. K. Morikawa, and K. Emoto. 2013. ’Compartmentalized calcium transients trigger dendrite pruning in Drosophila sensory neurons’, Science, 340: 1475–8.

Kawasaki, F., R. Felling, and R. W. Ordway. 2000. ’A temperature-sensitive paralytic mutant defines a primary synaptic calcium channel in Drosophila’, J Neurosci, 20: 4885–9.

Kittel, Robert J., Carolin Wichmann, Tobias M. Rasse, Wernher Fouquet, Manuela Schmidt, Andreas Schmid, Dhananjay A. Wagh, Christian Pawlu, Robert R. Kellner, Katrin I. Willig, Stefan W. Hell, Erich Buchner, Manfred Heckmann, and Stephan J. Sigrist. 2006. ’Bruchpilot Promotes Active Zone Assembly, Ca^2+^ Channel Clustering, and Vesicle Release’, Science, 312: 1051–54.

Kurdyak, P., H. L. Atwood, B. A. Stewart, and C. F. Wu. 1994. ’Differential physiology and morphology of motor axons to ventral longitudinal muscles in larval Drosophila’, J Comp Neurol, 350: 463–72.

Kurshan, P. T., A. Oztan, and T. L. Schwarz. 2009. ’Presynaptic alpha2delta-3 is required for synaptic morphogenesis independent of its Ca2+-channel functions’, Nat Neurosci, 12: 1415–23.

Laghaei, R., J. Ma, T. B. Tarr, A. E. Homan, L. Kelly, M. S. Tilvawala, B. S. Vuocolo, H. P. Rajasekaran, S. D. Meriney, and M. Dittrich. 2018. ’Transmitter release site organization can predict synaptic function at the neuromuscular junction’, J Neurophysiol, 119: 1340–55.

Li, Jingjun, James Ashley, Vivian Budnik, and Manzoor A. Bhat. 2007. ’Crucial Role of <EM<Drosophila>EM> Neurexin in Proper Active Zone Apposition to Postsynaptic Densities, Synaptic Growth, and Synaptic Transmission’, Neuron, 55: 741–55.

Li, Xiling, Pragya Goel, Joyce Wondolowski, Jeremy Paluch, and Dion Dickman. 2018. ’A Glutamate Homeostat Controls the Presynaptic Inhibition of Neurotransmitter Release’, Cell Reports, 23: 1716–27.

Lipscombe, D., A. Andrade, and S. E. Allen. 2013. ’Alternative splicing: functional diversity among voltage-gated calcium channels and behavioral consequences’, Biochim Biophys Acta, 1828: 1522–9.

Lipscombe, D., and E. J. Lopez Soto. 2019. ’Alternative splicing of neuronal genes: new mechanisms and new therapies’, Curr Opin Neurobiol, 57: 26–31.

Littleton, J. Troy, and Barry Ganetzky. 2000. ’Ion Channels and Synaptic Organization’, Neuron, 26: 35–43.

Liu, C., L. S. Bickford, R. G. Held, H. Nyitrai, T. C. Sudhof, and P. S. Kaeser. 2014. ’The active zone protein family ELKS supports Ca2+ influx at nerve terminals of inhibitory hippocampal neurons’, J Neurosci, 34: 12289–303.

Liu, K. S., M. Siebert, S. Mertel, E. Knoche, S. Wegener, C. Wichmann, T. Matkovic, K. Muhammad, H. Depner, C. Mettke, J. Buckers, S. W. Hell, M. Muller, G. W. Davis, D. Schmitz, and S. J. Sigrist. 2011. ’RIM-binding protein, a central part of the active zone, is essential for neurotransmitter release’, Science, 334: 1565–9.

Liu, S., P. Hoess, and J. Ries. 2022. ’Super-Resolution Microscopy for Structural Cell Biology’, Annu Rev Biophys, 51: 301–26.

Lnenicka, G. A., and H. Keshishian. 2000. ’Identified motor terminals in Drosophila larvae show distinct differences in morphology and physiology’, J Neurobiol, 43: 186–97.

Los, G. V., L. P. Encell, M. G. McDougall, D. D. Hartzell, N. Karassina, C. Zimprich, M. G. Wood, R. Learish, R. F. Ohana, M. Urh, D. Simpson, J. Mendez, K. Zimmerman, P. Otto, G. Vidugiris, J. Zhu, A. Darzins, D. H. Klaubert, R. F. Bulleit, and K. V. Wood. 2008. ’HaloTag: a novel protein labeling technology for cell imaging and protein analysis’, ACS Chem Biol, 3: 373–82.

Lu, Z., A. K. Chouhan, J. A. Borycz, Z. Lu, A. J. Rossano, K. L. Brain, Y. Zhou, I. A. Meinertzhagen, and G. T. Macleod. 2016. ’High-Probability Neurotransmitter Release Sites Represent an Energy-Efficient Design’, Curr Biol, 26: 2562–71.

Ly, Cindy V., Chi-Kuang Yao, Patrik Verstreken, Tomoko Ohyama, and Hugo J. Bellen 2008. ’straightjacket is required for the synaptic stabilization of cacophony, a voltage-gated calcium channel α1 subunit’, Journal of Cell Biology, 181: 157–70.

Macleod, G. T., L. Chen, S. Karunanithi, J. B. Peloquin, H. L. Atwood, J. E. McRory, G. W. Zamponi, and M. P. Charlton. 2006. ’The Drosophila cacts2 mutation reduces presynaptic Ca2+ entry and defines an important element in Cav2.1 channel inactivation’, Eur J Neurosci, 23: 3230–44.

McDonald, N. A., R. D. Fetter, and K. Shen. 2020. ’Assembly of synaptic active zones requires phase separation of scaffold molecules’, Nature, 588: 454–58.

Medeiros, Audrey T., and Kate O’Connor-Giles. 2023. ’To Ib or not to b: Transcriptional regulation of tonic type Ib vs. phasic type Is motor neurons’, Neuron, 111: 3497–99.

Melom, J. E., Y. Akbergenova, J. P. Gavornik, and J. T. Littleton. 2013. ’Spontaneous and evoked release are independently regulated at individual active zones’, J Neurosci, 33: 17253–63.

Miki, Takafumi, Walter A. Kaufmann, Gerardo Malagon, Laura Gomez, Katsuhiko Tabuchi, Masahiko Watanabe, Ryuichi Shigemoto, and Alain Marty. 2017. ’Numbers of presynaptic Ca^2+^ channel clusters match those of functionally defined vesicular docking sites in single central synapses’, Proceedings of the National Academy of Sciences, 114: E5246–E55.

Mrestani, Achmed, Martin Pauli, Philip Kollmannsberger, Felix Repp, Robert J. Kittel, Jens Eilers, Sören Doose, Markus Sauer, Anna-Leena Sirén, Manfred Heckmann, and Mila M. Paul. 2021. ’Active zone compaction correlates with presynaptic homeostatic potentiation’, Cell Reports, 37: 109770.

Muhammad, K., S. Reddy-Alla, J. H. Driller, D. Schreiner, U. Rey, M. A. Bohme, C. Hollmann, N. Ramesh, H. Depner, J. Lutzkendorf, T. Matkovic, T. Gotz, D. D. Bergeron, J. Schmoranzer, F. Goettfert, M. Holt, M. C. Wahl, S. W. Hell, P. Scheiffele, A. M. Walter, B. Loll, and S. J. Sigrist. 2015. ’Presynaptic spinophilin tunes neurexin signalling to control active zone architecture and function’, Nat Commun, 6: 8362.

Muller, C. S., A. Haupt, W. Bildl, J. Schindler, H. G. Knaus, M. Meissner, B. Rammner, J. Striessnig, V. Flockerzi, B. Fakler, and U. Schulte. 2010. ’Quantitative proteomics of the Cav2 channel nano-environments in the mammalian brain’, Proc Natl Acad Sci U S A, 107: 14950–7.

Nakamura, Yukihiro, Harumi Harada, Naomi Kamasawa, Ko Matsui, Jason S Rothman, Ryuichi Shigemoto, R. Angus Silver, David A DiGregorio, and Tomoyuki Takahashi. 2015. ’Nanoscale Distribution of Presynaptic Ca2+ Channels and Its Impact on Vesicular Release during Development’, Neuron, 85: 145–58.

Newman, Z. L., A. Hoagland, K. Aghi, K. Worden, S. L. Levy, J. H. Son, L. P. Lee, and E. Y. Isacoff. 2017. ’Input-Specific Plasticity and Homeostasis at the Drosophila Larval Neuromuscular Junction’, Neuron, 93: 1388–404 e10.

Newman, Zachary L., Dariya Bakshinskaya, Ryan Schultz, Samuel J. Kenny, Seonah Moon, Krisha Aghi, Cherise Stanley, Nadia Marnani, Rachel Li, Julia Bleier, Ke Xu, and Ehud Y. Isacoff. 2022. ’Determinants of synapse diversity revealed by super-resolution quantal transmission and active zone imaging’, Nature Communications, 13: 229.

Öztürk-Çolak, Arzu, Steven J Marygold, Giulia Antonazzo, Helen Attrill, Damien Goutte-Gattat, Victoria K Jenkins, Beverley B Matthews, Gillian Millburn, Gilberto dos Santos, Christopher J Tabone, and FlyBase Consortium. 2024. ’FlyBase: updates to the Drosophila genes and genomes database’, Genetics, 227.

Paez-Segala, M. G., M. G. Sun, G. Shtengel, S. Viswanathan, M. A. Baird, J. J. Macklin, R. Patel, J. R. Allen, E. S. Howe, G. Piszczek, H. F. Hess, M. W. Davidson, Y. Wang, and L. L. Looger. 2015. ’Fixation-resistant photoactivatable fluorescent proteins for CLEM’, Nat Methods, 12: 215–8, 4 p following 18.

Peled, E. S., and E. Y. Isacoff. 2011. ’Optical quantal analysis of synaptic transmission in wild-type and rab3-mutant Drosophila motor axons’, Nat Neurosci, 14: 519–26.

Peled, E. S., Z. L. Newman, and E. Y. Isacoff. 2014. ’Evoked and spontaneous transmission favored by distinct sets of synapses’, Curr Biol, 24: 484–93.

Peng, I. F., and C. F. Wu. 2007. ’Drosophila cacophony channels: a major mediator of neuronal Ca2+ currents and a trigger for K+ channel homeostatic regulation’, J Neurosci, 27: 1072–81.

Petersen, Sophie A., Richard D. Fetter, Jasprina N. Noordermeer, Corey S. Goodman, and Aaron DiAntonio. 1997. ’Genetic Analysis of Glutamate Receptors in Drosophila Reveals a Retrograde Signal Regulating Presynaptic Transmitter Release’, Neuron, 19: 1237–48.

Radulovic, T., W. Dong, R. O. Goral, C. I. Thomas, P. Veeraraghavan, M. S. Montesinos, D. Guerrero-Given, K. Goff, M. Lubbert, N. Kamasawa, T. Ohtsuka, and S. M. Young, Jr. 2020. ’Presynaptic development is controlled by the core active zone proteins CAST/ELKS’, J Physiol, 598: 2431–52.

Rebola, Nelson, Maria Reva, Tekla Kirizs, Miklos Szoboszlay, Andrea Lőrincz, Gael Moneron, Zoltan Nusser, and David A. Digregorio. 2019. ’Distinct Nanoscale Calcium Channel and Synaptic Vesicle Topographies Contribute to the Diversity of Synaptic Function’, Neuron, 104: 693–710.e9.

Reddy-Alla, S., M. A. Bohme, E. Reynolds, C. Beis, A. T. Grasskamp, M. M. Mampell, M. Maglione, M. Jusyte, U. Rey, H. Babikir, A. W. McCarthy, C. Quentin, T. Matkovic, D. D. Bergeron, Z. Mushtaq, F. Gottfert, D. Owald, T. Mielke, S. W. Hell, S. J. Sigrist, and A. M. Walter. 2017. ’Stable Positioning of Unc13 Restricts Synaptic Vesicle Fusion to Defined Release Sites to Promote Synchronous Neurotransmission’, Neuron, 95: 1350–64 e12.

Reuveny, Adriana, Marina Shnayder, Dana Lorber, Shuoshuo Wang, and Talila Volk. 2018. ’M-alpha2/delta promotes myonuclear positioning and association with the sarcoplasmic-reticulum’, Development, 145: dev159558.

Risher, W. Christopher, Namsoo Kim, Sehwon Koh, Ji-Eun Choi, Petar Mitev, Erin F. Spence, Louis-Jan Pilaz, Dongqing Wang, Guoping Feng, Debra L. Silver, Scott H. Soderling, Henry H. Yin, and Cagla Eroglu. 2018. ’Thrombospondin receptor α2δ-1 promotes synaptogenesis and spinogenesis via postsynaptic Rac1’, Journal of Cell Biology, 217: 3747–65.

Sauvola, Chad W., Yulia Akbergenova, Karen L. Cunningham, Nicole A. Aponte-Santiago, and J. Troy Littleton. 2021. ’The decoy SNARE Tomosyn sets tonic versus phasic release properties and is required for homeostatic synaptic plasticity’, eLife, 10: e72841.

Schöpf, Clemens L., Cornelia Ablinger, Stefanie M. Geisler, Ruslan I. Stanika, Marta Campiglio, Walter A. Kaufmann, Benedikt Nimmervoll, Bettina Schlick, Johannes Brockhaus, Markus Missler, Ryuichi Shigemoto, and Gerald J. Obermair. 2021. ’Presynaptic α2δ subunits are key organizers of glutamatergic synapses’, Proceedings of the National Academy of Sciences, 118: e1920827118.

Sheng, Jiansong, Liming He, Hongwei Zheng, Lei Xue, Fujun Luo, Wonchul Shin, Tao Sun, Thomas Kuner, David T. Yue, and Ling-Gang Wu. 2012. ’Calcium-channel number critically influences synaptic strength and plasticity at the active zone’, Nat Neurosci, 15: 998–1006.

Smith, L. A., X. Wang, A. A. Peixoto, E. K. Neumann, L. M. Hall, and J. C. Hall. 1996. ’A Drosophila calcium channel alpha1 subunit gene maps to a genetic locus associated with behavioral and visual defects’, J Neurosci, 16: 7868–79.

Thomas, G. D. 2000. ’Effect of dose, molecular size, and binding affinity on uptake of antibodies’,Methods Mol Med, 25: 115–32.

Tong, X. J., E. J. Lopez-Soto, L. Li, H. Liu, D. Nedelcu, D. Lipscombe, Z. Hu, and J. M. Kaplan. 2017. ’Retrograde Synaptic Inhibition Is Mediated by alpha-Neurexin Binding to the alpha2delta Subunits of N-Type Calcium Channels’, Neuron, 95: 326–40 e5.

Voigt, A., R. Freund, J. Heck, M. Missler, G. J. Obermair, U. Thomas, and M. Heine. 2016. ’Dynamic association of calcium channel subunits at the cellular membrane’, Neurophotonics, 3: 041809.

Wang, L. Y., and G. J. Augustine. 2014. ’Presynaptic nanodomains: a tale of two synapses’, Front Cell Neurosci, 8: 455.

Wang, T., R. T. Jones, J. M. Whippen, and G. W. Davis. 2016. ’alpha2delta-3 Is Required for Rapid Transsynaptic Homeostatic Signaling’, Cell Rep, 16: 2875–88.

Weiss, N., and G. W. Zamponi. 2017. ’Trafficking of neuronal calcium channels’, Neuronal Signal, 1: NS20160003.

Weyhersmuller, A., S. Hallermann, N. Wagner, and J. Eilers. 2011. ’Rapid active zone remodeling during synaptic plasticity’, J Neurosci, 31: 6041–52.

Yazaki, Junshi, Yusuke Kawashima, Taisaku Ogawa, Atsuo Kobayashi, Mayu Okoshi, Takashi Watanabe, Suguru Yoshida, Isao Kii, Shohei Egami, Masayuki Amagai, Takamitsu Hosoya, Katsuyuki Shiroguchi, and Osamu Ohara. 2019. ’HaloTag-based conjugation of proteins to barcoding-oligonucleotides’, Nucleic Acids Research, 48: e8–e8.

Zhang, Yanfeng, Ting Wang, Yimei Cai, Tao Cui, Michelle Kuah, Stefano Vicini, and Tingting Wang. 2023. ’Corrigendum: Role of α2δ-3 in regulating calcium channel localization at presynaptic active zones during homeostatic plasticity’, Frontiers in Molecular Neuroscience, 16.

